# FGF8 promotes lipid droplet accumulation via the FGFR1/p-p38 axis in chondrocytes

**DOI:** 10.1101/2025.04.15.648880

**Authors:** Minglei Huang, Haoran Chen, Jieya Wei, Caixia Pi, Mengmeng Duan, Xiaohua Pu, Zhixing Niu, Siqun Xu, Shasha Tu, Sijun Liu, Jiazhou Li, Li Zhang, Yang Liu, Hao Chen, Chunming Xu, Jing Xie

## Abstract

Chondrocytes store lipids in the form of lipid droplets (LDs) and maintain cartilage lipid metabolic homeostasis by consuming or regenerating LDs. This modulation is largely mediated by a series of biochemical factors. Fibroblast growth factor 8 (FGF8) is one of the most important factors involved in the proliferation, differentiation, and migration of chondrocytes, and has attracted increasing attention in the physiology and pathology of cartilage. However, the effect of FGF8 on LD accumulation in chondrocytes remains unclear. This study aimed to elucidate the role of FGF8 in LDs and explore the underlying biomechanism involved. The results showed that FGF8 promoted LD accumulation in chondrocytes by upregulating perilipin1 (Plin1) expression. FGF8 activates the cytoplasmic p-p38 signaling pathway via fibroblast growth factor receptor 1 (FGFR1) to increase LD accumulation in chondrocytes. Subsequent experiments with siRNAs and specific inhibitors further confirmed the importance of the FGFR1/p38 axis for LD accumulation in chondrocytes exposed to FGF8. These results increase our understanding of the role of FGF8 in the lipid metabolic homeostasis of chondrocytes and provide insights into the physiology and pathology of cartilage.

## 1. Introduction

The lipid droplet (LD), a principal hub organelle for intracellular lipid storage, consists of a neutral lipid core enveloped by a phospholipid monolayer membrane [1–3]. LD biogenesis begins mostly in the endoplasmic reticulum (ER), the main organelle involved in neutral lipid synthesis [4]. The first step in LD biosynthesis is the synthesis of neutral lipids such as triacylglycerols (TGs) and cholesterol esters (CEs) in the ER [4,5]. These neutral lipids are dispersed in the ER at low concentrations. With these continuous neutral lipids syntheses, biophysical processes result in the formation of neutral lipid lens (also called oil lens) [5]. A widely recognized model states that neutral lipids nucleate an oil phase that minimizes the entropy costs involved in disrupting the ER bilayer membrane (1,4,5). Subsequently, the oil phase transitions to the oil lens. The oil lens accommodates more neutral lipids and promote neutral lipids aggregation to form LDs [5]. After being released into the cytoplasm, these nascent LDs gradually develop into larger mature LDs by storing more lipids or merging each other [3–5]. In addition, LDs interact with other organelles such as mitochondria and lysosomes to regulate cellular metabolism and function. For example, Meng et al. reported that phosphofructokinase, a glycolytic enzyme, promotes LD-mitochondrion tethering to increase β-oxidation [6]. Menon et al. reported that ARL8B, a GTPase, promotes LD-lysosome contact and induces lysosomal lipolysis of LDs [7]. More importantly, lipophagy, an autophagy process that specifically targets LDs, plays an important role in maintaining the cellular energy supply and alleviating metabolic diseases such as atherosclerosis [8]. Lin et al. reported that the recovery of lipophagy via the inhibition of PPAR/PI3K/AKT signaling relieves atherosclerosis-related symptoms including lipid accumulation, apoptosis, and inflammation [9].

The primary function of intracellular LDs is to serve as storage reservoirs for neutral lipids and supply essential lipid precursors whenever the cellular lipid level decreases [10]. These neutral lipids can be used for cell energy supply through mitochondrial β-oxidation and for structural composition such as lipid membrane expansion [10]. In addition, LDs can assist cells in preventing lipid toxicity [11]. Free lipids such as fatty acids can act as detergents to disrupt the cell membrane structure. Synthesizing fatty acids into triglycerides and storing them in LDs effectively prevents intracellular lipid toxicity. However, despite the numerous positive effects of LDs within cells, excessive LD accumulation can result in the development of various diseases, such as hepatic steatosis [10,11]. As organelles with important functions in cells, LDs are also regulated by a variety of proteins [12]. These proteins, including a series of enzymes that regulate the dynamic process of LD biogenesis and linker proteins that connect LDs to other organelles, regulate LD activities [12]. These proteins are roughly divided into two types: integral and peripheral proteins. Integral proteins are inserted into the monolayer membrane of LDs and can recruit peripheral proteins to the LDs [12]. Integral proteins play vital roles in the generation, maintenance, and lipolysis of LDs, and are composed of two functional subclasses according to their trafficking pathways. One is Class I LD protein that originates in the ER [12]. These proteins are then transferred to LDs during LD formation. For example, seipin, a typical ER protein located at the site of LD budding, can form a flexible cage to promote LD formation [13]. Furthermore, it plays a pivotal role in facilitating the connection between nascent LDs and the ER, and enhancing the transfer of lipids and proteins from the ER to LDs [14]. In addition to seipin, some lipid synthase- and ubiquitination-related proteins, such as diacylglycerol acyltransferase 2 (DGAT2) and ancient ubiquitous protein 1 (AUP1) [15–17], also belong to this subclass. Malis et al. reported that DGAT2 is transferred from the ER to the LD surface and promotes LD growth [17]. Smith et al. reported that AUP1 can reduce the formation of misfolded proteins and is thus important for protein quality control in the progression of LD formation and maturation [18]. Moreover, a decrease in AUP1 expression can reduce the accumulation of cellular lipids and affect the final formation of LDs [15,16,19]. The other is Class II LD protein, which is synthesized in the cytoplasmic ribosome and targets LDs. The typical protein is the perilipin (Plin) protein family, which contains five family members (Plin1-5) [12,20]. The main function of Plins is to inhibit the lipolysis of LDs, but in the case of an urgent need for a large number of liposomes, Plins can be phosphorylated with the help of protein kinase A (PKA), and the phosphorylated Plins then recruit lipases such as adipose triglyceride lipase (ATGL) and hormone sensitive lipase (HSL) to achieve lipolysis. In addition, Plins play a role in the interactions between LDs and other organelles. For example, Miner et al reported that phosphorylated Plin5 interacts with fatty acid transport protein 4 (FATP4) to promote the interaction between LDs and the mitochondria, which facilitates fatty acid transport from LDs to the mitochondria [21].

Cartilage tissue is characterized by the absence of blood vessels, lymphatic vessels, and nerves [22,23]. Owing to its unique composition, which is composed mainly of type II collagen and proteoglycans, cartilage tissue has superior performance in resistance to compression and decompression, but also has extremely poor healing ability once injury or damage occurs [24,25]. Cartilage injury or damage can lead to osteoarthritis (OA), a typical degenerative disease [26]. Recent studies have also revealed that OA pathogenesis is highly similar to that of systemic metabolic syndrome (MS), with lipid metabolism disorder as a vital indicator [27–29]. For example, Park et al. reported that excessive LD accumulation in chondrocytes induced by PPARα or ACOT12 deficiency accelerated the degradation of the cartilage matrix in OA and suggested that a reduction in LD accumulation in chondrocytes may be a potential therapeutic target for OA [30]. Wang et al. reported that GDF11 inhibits abnormal lipid formation and LD accumulation by promoting ubiquitination of PPARγ in chondrocytes in temporomandibular joint osteoarthritis (TMJOA) [31]. Chondrocytes can obtain nutrients only from the subchondral bone and the surrounding synovial fluid, indicating that chondrocytes need to store sufficient raw lipid materials as energy reserves [32,33]. These raw lipids stored in cellular LDs include fatty acids, triacylglycerols, cholesterol esters, and their byproducts, including prostaglandins and leukotrienes [34]. Healthy chondrocytes have high levels of omega-9 fatty acids (n-9 FAs) and low levels of omega-6 polyunsaturated fatty acids (n-6 PUFAs) [35]. With increasing chondrocyte age, the content of n-6 PUFAs increases, whereas that of n-9 FAs decreases. In addition, arachidonic acid (a typical inflammation-related lipid) and n-6 PUFA are also increased in osteoarthritic chondrocytes compared to healthy chondrocytes [36]. After binding to glycerol, triacylglycerol is the main intracellular storage form of fatty acids. When chondrocytes urgently require fatty acids, triacylglycerols are gradually broken down into fatty acids and glycerol by a variety of catabolic enzymes to provide the raw material supply for mitochondrial aerobic respiration [1,35]. Cholesterol esters also affect the growth, physiological activity, and differentiation of chondrocytes [35]. When the regulatory proteins that assist in the transport of cholesterol esters out of chondrocytes are inhibited, cholesterol esters accumulate in chondrocytes, leading to sharp deterioration of the OA cartilage matrix [37]. In addition, prostaglandins and leukotrienes have been recognized for their roles in promoting inflammatory responses in cartilage joints, although their proportion in lipids is much lower. Both prostaglandins and leukotrienes are biosynthesized from arachidonic acid using prostaglandin synthase and lipoxygenase, respectively [38,39]. Prostaglandin E2 (PGE2), a typical prostaglandin, has been implicated in the promotion of chondrocyte hypertrophy. Interestingly, a recent study revealed an unexpected role for PGE2, showing that it inhibited the production of related hypertrophy signals in hypertrophic chondrocytes [40]. Leukotriene B4 (LTB4), a potent proinflammatory leukotriene synthesized by leukotriene A_4_ hydrolase (LTA_4_H), has been shown to be associated with cartilage matrix degeneration [41].

LD formation and maturation are mediated by a series of biochemical factors, including interleukins, heparan sulfates, and transforming growth factors during OA progression [42–43]. FGF8, an important member of the fibroblast growth factor family, plays a vital role in chondrocyte proliferation, differentiation, and extracellular matrix synthesis during the early stages of vertebrate development [44–45]. In cartilage diseases, FGF8 has been implicated in the pathogenesis of Kashin-Beck disease, where it disrupts chondrogenic differentiation through FGFR3 and enhances chondrocyte proliferation and hypertrophy [46]. FGF8 has been reported to promote cartilage degradation and exacerbate OA by increasing the production of matrix metalloproteinase-3 [37,44,47]. We recently confirmed that FGF8 regulates the expression of gelatinases (Mmp-2 & -9) and thus, has a direct effect on the degradation of the cartilage matrix [48]. However, whether and how FGF8 participates in lipid metabolism in chondrocytes during OA progression remains unclear. In this study, we aimed to investigate the role of FGF8 in the accumulation of LDs in chondrocytes and to explore the underlying biomechanisms. In this study, we aimed to increase the understanding of the lipid accumulation in chondrocytes and provide potential strategies for the prevention and treatment of osteoarticular diseases.

## Materials and Methods

### 1. Chondrocyte acquisition and culture

All experiments involving animal samples (mouse samples) were conducted in accordance with the protocols and ethical principles approved by the Institutional Review Board at the West China Hospital of Stomatology (No. WCHSIRB-CT-2022–127). Primary chondrocytes were obtained as previously described [49,50]. First, articular cartilage was isolated from newborn C57BL/6J mice (2-3 days old) and immersed in phosphate buffer solution (PBS) containing 1% penicillin-streptomycin (SV30010, Hyclone, Logan, UT, USA). Then, the articular cartilage was digested with 0.25% trypsin (25200056, Thermo Fisher Scientific, Waltham, MA, USA) for 30 min at 37°C and then treated with 0.5% type II collagenase (C6885, Sigma-Aldrich, St. Louis., MO, USA) dissolved in Dulbecco’s Modified Eagle’s Medium (DMEM) (SH30243.01B, HyClone, Logan, UT, USA) for 12 h at 37°C. Next, the digested solution was neutralized with DMEM supplemented with 10% fetal bovine serum (FBS) (SH30406.06, Hyclone, Logan, UT, USA), and centrifuged at 1,000 rpm for 8 min. The supernatant was discarded, and the chondrocytes in the pellet were resuspended in fresh DMEM containing 1% penicillin-streptomycin and 10% FBS. Chondrocytes were then incubated at 37°C in a humidified atmosphere with 5% CO_2_ and the culture medium was changed every two days. The first three generations of chondrocytes were used for subsequent experiments.

### 2. Transfection of small interfering RNA (siRNA)

Chondrocytes were cultured in antibiotic-free 10% FBS DMEM for 24 h, and then transfected with siRNA at a final concentration of 50 nM according to the manufacturer’s instructions. The transfection reagent used in this study was Lipofectamine RNAiMAX (13778075, Invitrogen, Burlington, ON, Canada). Chondrocytes transfected with si-NC were used as negative controls. The siRNA (hanbio, Shanghai, CN) sequences were as follows: si-NC: sense, 5’-UUCUCCGAACGUGUCACGUTT-3’, anti-sense, 5’-ACGUGACACGUUCGGAGAATT-3’; si-Plin1: sense,5’-GGCUCUGUCAUCAUCUAUATT-3’, anti-sense, 5’-UAUAGAUGAUGACAGAGCCTT-3’; and si-FGFR1: sense, 5’-AGAUCCCGGCUCUUCAAUATT-3’, anti-sense, 5’-UAUUGAAGAGCCGGGAUCUTT-3’. Knock-down efficiency was verified using qPCR.

### 3. RNA extraction and quantitative real-time polymerase chain reaction (qPCR)

Chondrocytes were treated with FGF8 at 25 ng/ml for 12 h before RNA extraction. Total RNA was extracted using RNA Isolation Kit (RE-03011, FOREGENE, Chengdu, CN). After collection, the total RNA was reverse-transcribed into complementary DNA (cDNA) using a reverse transcription kit (K1621-RevertAid, Mbi, MD). qPCR was performed using the SYBR Premix Ex II Taq PCR kit (RR820A, Takara, Tokyo, JPN) in a real-time PCR detection system (CFX 96, Bio-rad, JPN). Primer sequences used in this study included β-actin (Forward: 5’-GGCTGTATTCCCCTCCATCG-3’, Reverse: 5’-CCAGTTGGTAACAATGCCATGT-3’), FGFR1 (Forward: 5’-GCTTCATCTACGGAATGTCTCC-3’, Reverse: 5’-TCTTCCAGGGCGATAGAGTTAC-3’), and Plin1 (Forward: 5’-GACTGAGGTGGCGGTCTGCTGC-3’, Reverse: 5’-GGGGTGGGCTTCTTTGGTGCTG-3’). The 2^−ΔΔCt^ method was used to analyze results. β-actin was used as an internal control.

### 4. RNA sequencing

Total RNA was extracted from chondrocytes treated with FGF8 at 25 ng/ml for 24 h using Trizol (15596-018, Thermo Fisher Scientific, Waltham, MA, USA) and sent to Shanghai Lifegenes for transcriptome sequencing analysis as described in previous articles [51]. RNA integrity was assessed using the RNA Nano 6000 Assay Kit of the Bioanalyzer 2100 system. The RNA input was set to be 1.5 μg per sample. Index-coded samples were categorized using the HiSeq 4000 PE Cluster Kit, according to the manufacturer’s instructions. Read numbers of the genes were calculated using HTSeq v0.6.1, followed by fragments per kilobase of exon model per million mapped fragments (FPKM) calculation. A Pheatmap was generated using the R programming language. GO terms with a p-value less than 0.05 were considered significantly enriched by differentially expressed genes. Kyoto Encyclopedia of Genes and Genomes (KEGG) functional enrichment analysis was performed using KOBAS v3.0 software.

### 5. Western blotting

The detailed procedure of western blotting was followed as previously described [52]. Chondrocytes were cultured in DMEM containing 10% FBS until they reached to an appropriate density (80∼90% confluency). The culture medium was then replaced with DMEM containing 2% FBS to starve chondrocytes for 12 h. Next, chondrocytes were treated with FGF8 (100-25A, PeproTech, 100-25, USA) at 5, 10, or 25 ng/ml in the presence or absence of SB203580 at 20 µM (MCE, HY-10256, USA) in 1% FBS DMEM. At the indicated timepoints, the culture medium was discarded, and chondrocytes were lysed using RIPA lysis solution (P0013B, Beyotime, Shanghai, CN) containing 1% PMSF (P7626, sigma, GER) on an ice bath. The lysates were mixed with an equal volume of loading buffer (1610737EDU, Bio-Rad, Hercules, CA, USA) and boiled at 100°C for 5-10 min to prepare protein samples. The protein samples were separated by electrophoresis using 10% sodium dodecyl sulfate-polyacrylamide gels (1.0 mm) and transferred to PVDF membranes (IPVH00010, Millipore Billerica, MA, USA). The PVDF membranes were blocked with 5% skim milk for 1-2 h and incubated with primary antibodies overnight at 4°C. The primary antibodies used in this study were as follows: β-actin (200068-8F10, 1:1000, anti-mouse, Zenbio, CN), Erk (343830, 1:1000, anti-mouse, Zenbio, CN), JNK (R22866, 1:1000, anti-mouse, Zenbio, CN), p38 (R25239, 1:1000, anti-mouse, Zenbio, CN), p-p38 (310091, 1:1000, anti-mouse, Zenbio, CN), p-Erk (310065, 1:1000, anti-mouse, Zenbio, CN), p-JNK (381100, 1:1000, anti-mouse, Zenbio, CN), β-catenin (201328-5D6, 1:1000, anti-mouse, Zenbio, CN). The PVDF membranes were then washed three times for 5 min using Tris-buffered saline with 0.05% Tween 20 (TBST). Next, the PVDF membranes were incubated with the corresponding secondary antibodies (511203 and 511103, 1:5000, Zenbio, CN) for 2-3 h. Finally, signals on the PVDF membranes were detected using the Immobilon^®^ Western Kit (P90719, Millipore, Billerica, MA, USA). The signal results were analyzed using ImageJ software.

### 6. Immunofluorescence staining and BODIPY493/503 staining assay

The detailed immunofluorescent procedure was followed as previously described [53]. Briefly, chondrocytes were washed three times with 1×PBS, fixed with 4% paraformaldehyde for 15 min, and permeabilized with 0.25% Triton X-100 (P0096, Beyotime, Shanghai, CN) for 10 min. After permeabilization, chondrocytes were blocked with 5% BSA (V900933, Sigma-Aldrich, MO, USA) for 2 h and incubated with primary antibodies (Plin1: 1:200, CST (3467), USA; p-p38: 1:200, Zenbio (310091), CN) overnight at 4 °C. After primary antibodies incubation, the samples were incubated with the corresponding secondary antibody (Alexa Fluor 647 donkey anti-rabbit IgG, 1:200, ab150075, Abcam, UK) for 2 h. For LD staining, chondrocytes were incubated with BODIPY493/503 dye (D3922, Thermo Fisher Scientific, Waltham, MA, USA) for 10 min. Next, chondrocyte cytoskeletons were stained with FITC (A12379, R415, Invitrogen, Carlsbad, CA, USA) at 4 °C overnight. The chondrocyte nuclei were then stained with DAPI (D9542, Sigma, St. Louis, MO, USA) for 10 min. Immunofluorescence images were captured using the confocal laser scanning microscope (CLSM, FV3000, Olympus, Japan) and analyzed by ImageJ.

### 7. AAV-FGF8 animal model and immunofluorescence staining

FGF8 overexpression in articular cartilage was achieved by injecting of adeno-associated virus (AAV) carrying the FGF8 gene, as previously described [50]. AAV-FGF8 was obtained from GENECHEM (Shanghai, CN). The carriers used are CMV bGlobin-MCS-EGFP-3FLAG-WPRE-hGH polyA. The gene ID was 14179 and transcript ID was NM_-_010205 in NCBI. Briefly, after enzyme digesting the vector, the PCR-amplified target fragment is ligated to the vector. And then, the ligated vector was transfected into DH5α sensory cells. After transfecting, the DH5α sensory cells were cultured for 12-16 h and the positive clones were screened for PCR identification. After PCR identification, the positive DH5α sensory cells were subjected to expansion culture and plasmid extraction. After constructing the plasmid carrying FGF8, the constructed vector plasmid was transfected into 293T cells (packaging cells for AAV, adherent-dependent epithelioid cells) by the triple plasmid transfection system (pAAV-RC plasmid, pHelper plasmid, shuttle plasmid). After transfecting with plasmid carrying FGF8, cell precipitates were collected 72 h for virus purification. AAV-FGF8 was administered into the temporomandibular joint cavity under anaesthetic conditions (5 × 10^10^ V.g/ml, 3 μl per joint). Temporomandibular cartilage samples were harvested after 4 weeks’ injection. Immunofluorescence analysis was performed using antibodies or liquid kit (BODIPY493/503). Briefly, Tissue samples were incubated with primary antibodies (osterix, 1:200, ab209484, Abcam, UK; Plin1: 1:200, 3467, CST, USA) overnight at 4 °C. After incubation with the primary antibodies, the corresponding secondary antibodies (Alexa Fluor 647 donkey anti-rabbit IgG, 1:200, ab150075, Abcam, UK; Alexa Fluor 488 Goat Anti-Mouse IgG, 1:200, ab150133, Abcam, UK) were used and incubated for 2 h. After each antibody incubation, the samples were washed 6 times with PBST (containing 0.05% Tween 20) for 10 min. For liquid staining, the BODIPY493/503 dye was incubated overnight at 4 °C. Nuclear staining was performed using DAPI for 10 min. The images were obtained by CLSM.

### 8. Transfection of lentivirus

Chondrocytes were cultured to 30%-50% confluence in 10% FBS DMEM, and then transfected with lentivirus carrying Plin1 (hanbio, Shanghai, CN) at a final concentration of 30 MOI (multiplicity of infection) according to the manufacturer’s instructions. The carrier of the Plin1 gene was pHBLV-CMV-MCS-3FLAG-EF1-ZsGreen-T2A-PURO. The gene and lentivirus information is shown in Table S3. We use the one-half infection method for infection. Briefly, only half of the fresh complete medium was added at the time of lentivirus infection, and the medium was replenished 4 h later. When we performed lentivirus transfection, we also used 5 µg/ml polybrene (hanbio, Shanghai, CN) to aid transfection according to the manufacturer’s instructions. The Chondrocytes were infected for 72 h before testing. We used GFP tracing by CLSM and western blotting to measure transfection efficiency. In addition, due to the 488 nm GFP carried by the lentivirus, we used another neutral lipid droplet stain kit, HCS LipidTOX^TM^ Deep Red neutral lipid Stain (H34477, Thermo Fisher Scientific, Waltham, MA, USA), to observe neutral LDs.

### 9. Statistical analysis

The results were based on three replications and analyzed using GraphPad Prism 9.0.0 software (GraphPad Software Inc., San Diego, CA, USA). Two-tailed Student’s t test for two groups and one-way analysis of variance for multiple groups were used to test differences. And differences were considered statistically significant at p < 0.05.

## Results

### 1. FGF8 promotes lipid droplet accumulation in chondrocytes

To investigate the effect of FGF8 on chondrocyte metabolism, we performed RNA sequencing on chondrocytes exposed to FGF8 at 25 ng/ml for 24 h. We clustered all the altered pathways via Kyoto Encyclopedia of Genes and Genomes (KEGG) analysis and found multiple lipid metabolism-related pathways, such as fat absorption and digestion, ether lipid metabolism, glycosphingolipid biosynthesis, regulation of lipolysis in adipocytes, PPAR signaling pathway, and arachidonic acid metabolism pathways, were upregulated (**Figure 1A**, red boxes). These upregulated signaling pathways are tightly correlated with lipid abundance and distribution [54,55] and indicate potential changes in LDs in chondrocytes after FGF8 treatment. To visualize the changes in neutral lipid and LD accumulation in chondrocytes, we performed fluorescence staining using BODIPY493/503 in chondrocytes exposed to FGF8 at 25, 50, and 100 ng/ml (**Figure 1B** at high magnification **& S1** at low magnification), and found that FGF8 significantly increased neutral lipid accumulation by increasing the number of LDs formed. Total fluorescence quantification confirmed the increase in the number of neutral LDs in the chondrocytes (**Figure 1C**). In addition, quantification of the number of visible LDs further confirmed that FGF8 increased the accumulation of neutral lipids and LDs in the chondrocytes (**Figure 1D**). By injecting an adeno-associated virus (AAV) carrying the FGF8 gene into the mouse temporomandibular joint to achieve FGF8 overexpression in cartilage tissue, we found that neutral lipid and LD accumulation was largely enhanced (**Figure 1E**). We observed that neutral lipid accumulation was increased in both the proliferative layer (white arrows, boxed area) and the hypertrophic layer (yellow arrows, boxed area). By quantifying the neutral lipids in the entire cartilage layer, we confirmed the enhancement of neutral lipid accumulation in cartilage induced by FGF8 (**Figure 1F**).

**Figure 1.**
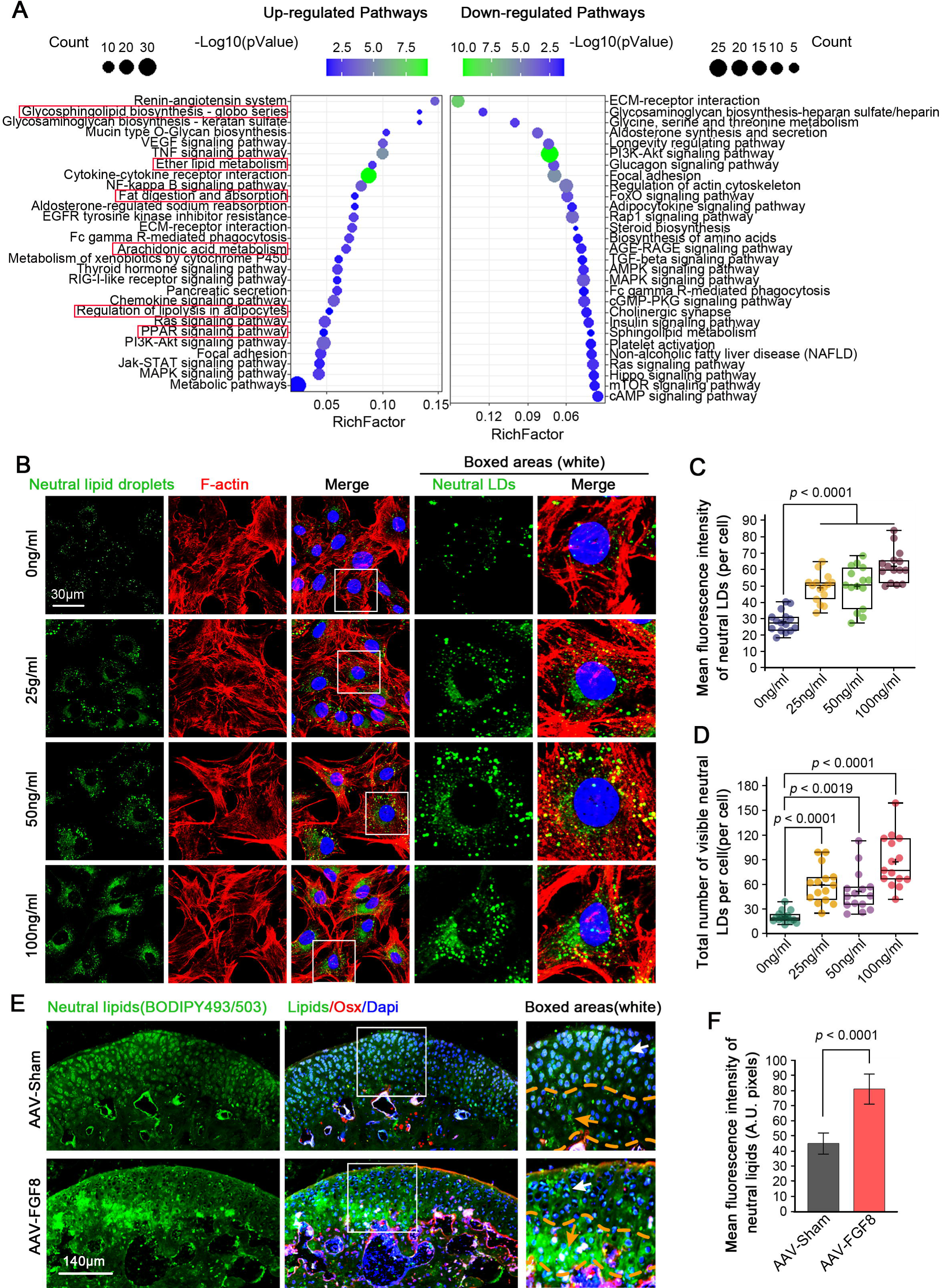
FGF8 induces lipid droplet accumulation in chondrocytes. **(A)** KEGG analysis based on RNA sequencing data showing changes in signaling pathways in chondrocytes exposed to FGF8 at 25 ng/ml. The left side indicated upregulated pathways, and the right side indicated downregulated pathways. The red boxes referred to the signaling pathways closely associated with lipid droplet accumulation. **(B)** Representative fluorescence images showing changes in lipid droplet**s** in chondrocytes exposed to different concentrations of FGF8. Chondrocytes were treated with FGF8 at 0, 25, 50, and 100 ng/ml for 2 days and then imaged using a 60 × CLSM. Green fluorescence indicated lipids, red fluorescence indicated the cytoskeleton (F-actin), and blue fluorescence indicated the nuclei. The white dashed boxes indicated the magnified regions. **(C)** Total fluorescence quantification per cell confirming changes in lipid accumulation in chondrocytes treated with 0, 25, 50, or 100 ng/ml FGF8 for 2 days. The data were based on 15 cells from three independent samples (n = 3). **(D)** Quantification of the number of visible lipid droplets per cell, confirming the changes in the number of lipid droplets in chondrocytes exposed to 0, 25, 50, and 100 ng/ml FGF8 for two days. The data were based on 15 cells from three independent samples (n = 3). **(E)** Representative fluorescence images showing changes in lipid accumulation in cartilage caused by AAV-FGF8 overexpression. Green fluorescence indicated lipids, red fluorescence indicated Osx staining (negative in mature cartilage and positive in calcified cartilage), and blue fluorescence indicated the nuclei. The white dashed boxes indicated the magnified regions. **(F)** Quantification of the mean fluorescence intensity confirming the changes in lipid accumulation in the cartilage exposed to FGF8. The data were based on three replications (n = 3). The data in **C, D** and **F** were based on a one-way analysis of variance. Data in **C** and **D** were shown in box (from 25%, 50% to 75%) and whisker (minimum to maximum) plots. Differences were considered statistically significant at p < 0.05.

### 2. FGF8 promotes lipid droplet accumulation by upregulating perilipin1 (Plin1) in chondrocytes

To explore how FGF8 promotes LD accumulation in chondrocytes, we clustered differentially expressed gene candidates that were tightly correlated with LD maturation based on RNA sequencing in the form of a pheatmap (**Figure 2A**). Through protein-protein interaction analysis using STRING (**Figure S3**), we established an interaction network among these candidates for FGF8-mediated LD accumulation. Notably, we found that the expression of Plin1, which plays a major role in LD formation and maturation by enveloping LDs to protect them against breakdown [20], was significantly increased in chondrocytes exposed to 25 ng/ml FGF8 (red box). We then performed qPCR to confirm the gene change in Plin1 in chondrocytes exposed to 25 ng/ml FGF8, and found that FGF8 increased the gene expression of Plin1 up to 1.85-fold relative to that in the control group (**Figure 2B**). At the protein level, western blotting showed that the expression of Plin1 increased in chondrocytes induced by FGF8 (**Figure 2C**), and quantitative analysis of Plin1 protein confirmed this result (**Figure 2D**). We also detected protein changes in hormone sensitive lipase (HSL), diacylglycerol O-acyltransferase 2 (DGAT2), and adipose triglyceride lipase (ATGL) in chondrocytes induced by FGF8 (**Figure S4**), and found that HSL was significantly decreased. Next, we performed immunofluorescence analysis to explore the protein expression and distribution of Plin1 in chondrocytes exposed to FGF8 (**Figure 2E**). The results revealed that Plin1 was mainly distributed in the cytoplasm around the nucleus and its expression increased in chondrocytes in response to 25 ng/ml FGF8. Using total fluorescence quantification, we further confirmed the increase in Plin1 expression in the chondrocytes (**Figure 2F**). In addition, in the cartilage layer, we detected increased Plin1 expression induced by AAV-FGF8 overexpression (**Figure S2**). RNA interference was performed to verify the importance of Plin1 in FGF8-mediated LD accumulation. After confirming the knockdown efficiency of si-Plin1 at 50 nM (**Figure 2G**), we performed immunofluorescence (**Figure 2H** at high magnification **& S5** at low magnification), and the results showed that knockdown of Plin1 clearly reduced LD accumulation in chondrocytes exposed to FGF8. Total fluorescence quantification further confirmed the reduced protein expression of Plin1 in chondrocytes caused by RNA interference in the presence of FGF8 (**Figure 2I**). Quantification of the visible LD number (**Figure 2J**) and total fluorescence per cell (**Figure S6**) confirmed the changes in LDs in chondrocytes induced by si-Plin1 in the presence of FGF8. Furthermore, we overexpressed Plin1 using lentivirus transfection to show the effect of Plin1 overexpression on LD accumulation. First, we used CLSM to detect the transfection efficiency of Plin1 overexpression in chondrocytes at 30 MOI and found that the lentivirus carrying Plin1 was successfully transfected into chondrocytes (**Figure S7**). At the protein level, we performed immunofluorescence to confirm the overexpression of Plin1 in chondrocytes after lentivirus transfection (**Figure 2L**), and total fluorescence quantification verified the increase in Plin1 protein in chondrocytes (**Figure 2M**). Using western blotting, we confirmed that the expression of Plin1 was increased (close to 2-fold high) in chondrocytes after Plin1 overexpression relative to the vector control **(Figure 2N &2O)**. After confirming the overexpression efficiency of LVs, we next investigated LD accumulation in chondrocytes after Plin1 overexpression (**Figure 2P & S8**). The result showed that the overexpression of Plin1 significantly promoted LD accumulation in chondrocytes. Quantification of the visible LD number (**Figure 2Q**) and total fluorescence of lipids per cell (**Figure S9**) confirmed that these changes in chondrocytes were induced by Plin1 overexpression. Taken together, these results demonstrate that FGF8 enhances LD accumulation by upregulating the expression of Plin1 in chondrocytes (**Figure 2K**).

**Figure 2.**
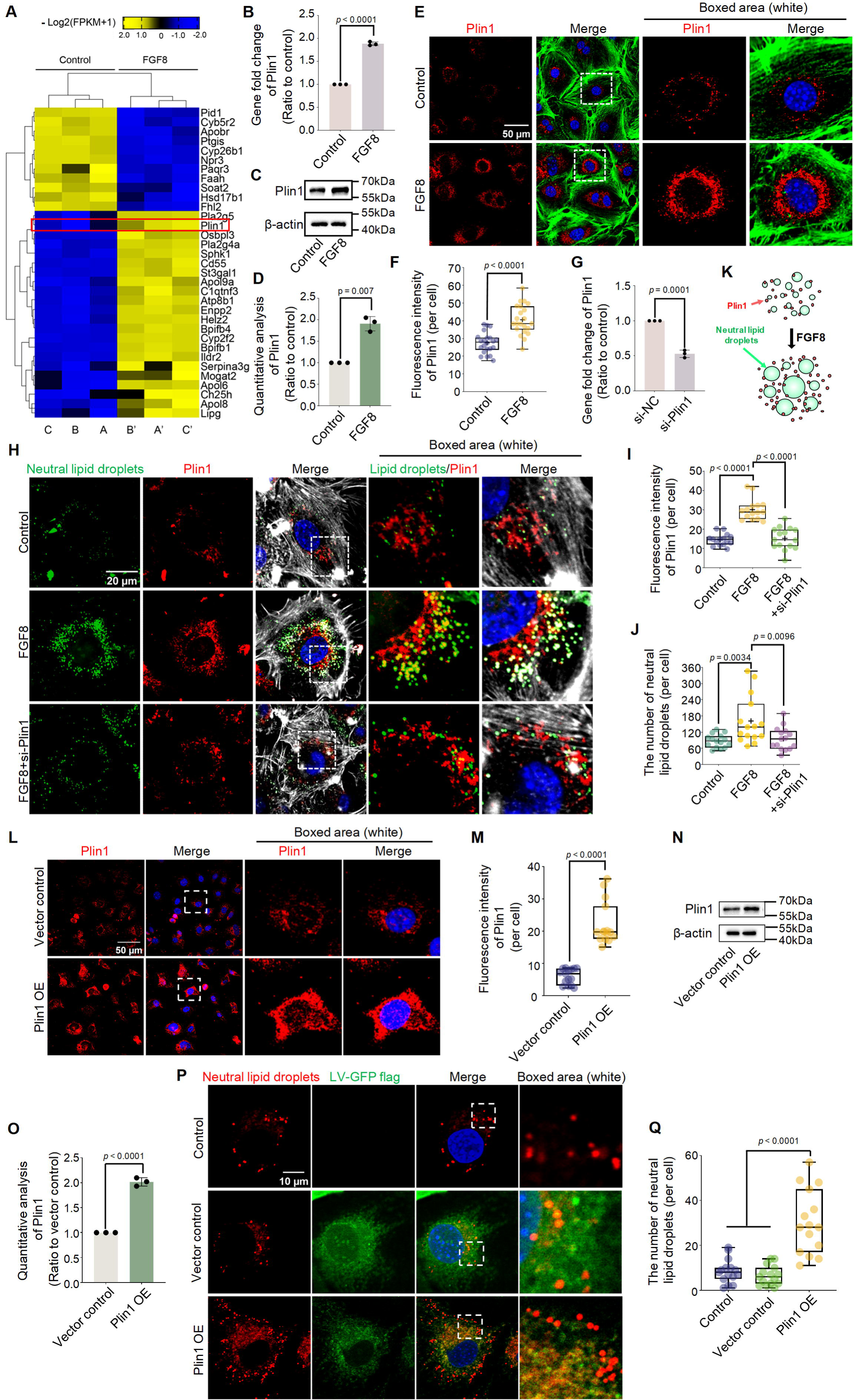
FGF8 enhances lipid droplet accumulation by upregulating Plin1 in chondrocytes. **(A)** The pheatmap based on RNA sequencing indicating changes in the levels of candidate mediators involved in lipid droplet accumulation. The red box indicated the gene changes in Plin1. Chondrocyte samples A & A’, B & B’, and C & C’ were independent of from the same parental chondrocytes. The gene candidates were presented as –log2(1+FPKM) and generated using the online R package, FPKM, Fragments Per Kilobase of exon model per Million mapped fragments. **(B)** qPCR verifying the change in Plin1 expression in chondrocytes exposed to FGF8 at 25 ng/ml. The data were based on three replications (n = 3). **(C)** Western blotting showing upregulation of Plin1 in chondrocytes induced by FGF8. β-actin was used as the internal reference. Data were representative of three independent samples (n = 3). **(D)** Quantitative analysis confirming the fold change in Plin1 protein expression in chondrocytes treated with FGF8 in (C). The data were based on three replicates (n = 3). **(E)** Representative fluorescence images showing changes in Plin1 in chondrocytes exposed to 25 ng/ml FGF8. Fluorescence images were captured using CLSM at 60×magnification. Red fluorescence indicated Plin1, green fluorescence indicated the cytoskeleton (F-actin), and blue fluorescence indicated the nuclei. The white dashed boxes indicated the magnified regions. **(F)** Total fluorescence quantification per cell confirming the changes in Plin1 protein level in (E) in chondrocytes treated with FGF8 at 25 ng/ml. The data were based on 21 cells from three independent samples (n = 3). **(G)** qPCR verifying the knockdown efficiency of 50 nM si-Plin1 in chondrocytes. The data were based on three replicates (n = 3). **(H)** Representative fluorescence images of individual cells indicating changes in lipid droplets in chondrocytes exposed to si-Plin1 in the presence of FGF8 at 25 ng/ml. Chondrocytes were pretreated with siRNA for 12 h and then treated with FGF8 at 25 ng/ml for 2 days. Fluorescence images were obtained using 60 × CLSM. Green fluorescence indicated lipid droplets, red fluorescence indicated Plin1, gray fluorescence indicated the cytoskeleton (F-actin), and blue fluorescenceindicated the nuclei. The white dashed boxes indicated the magnified regions. **(I)** Total fluorescence quantification per cell confirming the changes in Plin1 protein in (F) chondrocytes exposed to si-Plin1 in the presence of FGF8. The data were based on 15 cells from three independent samples (n = 3). **(J)** Quantification of the number of visible lipid droplets per cell confirming the changes in the number of lipid droplets in chondrocytes inhibited by si-Plin1 in the presence of FGF8 at 25 ng/ml. The data were based on 15 cells from three independent samples (n = 3). **(K)** Schematic diagram showing that Plin1 protects against the breakdown of lipid droplets and promotes the formation of larger lipid droplets in the presence of FGF8. **(L)** Representative fluorescence images showing change in Plin1 in chondrocytes after Plin1 overexpression by lentivirus. Fluorescence images were obtained using 60 × CLSM. Red fluorescence indicated Plin1, and blue fluorescence indicated the nuclei. The white dashed boxes indicated the magnified regions. **(M)** Total fluorescence quantification per cell confirming change in Plin1 protein level in chondrocytes in (L). The data were based on 15 cells from three independent samples (n = 3). **(N)** Western blotting showing increased expression of Plin1 in chondrocytes after Plin1 overexpression by lentivirus. β-actin was used as the internal reference. Data were representative of three independent samples (n = 3). **(O)** Quantitative analysis confirming the fold change of Plin1 protein in chondrocytes in (N). The data were based on three independent samples (n = 3). **(P)** Representative fluorescence images of individual cells showing change in lipid droplets in chondrocytes after Plin1 overexpression by lentivirus. Fluorescence images were obtained using 60 × CLSM. Green fluorescence indicated GFP protein carried by lentivirus, red fluorescence indicated lipid droplets, and blue fluorescence indicated the nuclei. The white dashed boxes indicated the magnified regions. **(Q)** Quantification of the number of visible lipid droplets per cell confirming the changes in the lipid droplet accumulation in chondrocytes in (P). The data were based on 15 cells from three independent samples (n = 3). The data in **F**, **I**, **J, M, N,** and **Q** were analyzed using one-way analysis of variance. The data in **B**, **D**, **G**, **M**, **O**, and **Q** were based on two-tailed Student’s t-tests. Data in **B**, **D**, **G**, and **O** were presented as the means ±SDs. The data in **F**, **I**, **M** and **Q** were shown in the box (from 25%, 50% to 75%) and whisker (minimum to maximum) plots. Differences were considered statistically significant at p < 0.05.

### 3. FGF8-mediated lipid droplet accumulation requires the participation of FGF receptor 1 (FGFR1)

To explore how FGF8 enters chondrocytes to initiate downstream responses, we clustered the differentially expressed FGF receptors via RNA sequencing and found that only FGFR1 was upregulated in chondrocytes treated with 25 ng/ml FGF8 (**Figures 3A and S10**). We then performed qPCR to confirm the increased gene expression of FGFR1 in chondrocytes (**Figure 3B**). To determine the role of FGFR1 in FGF8-mediated LD accumulation in chondrocytes, we used siRNAtargeting FGFR1. After confirming the knockdown efficiency of si-FGFR1 at 50 nM (**Figure 3C**), we detected the gene expression of Plin1 via qPCR (**Figure 3D**), and the results revealed that si-FGFR1 strongly decreased the increase in Plin1 gene expression in chondrocytes exposed to FGF8. Using immunofluorescence, we observed that si-FGFR1 effectively impaired the increase in cytoplasmic Plin1 protein levels in chondrocytes exposed to FGF8 (**Figure 3E**). Fluorescence quantification confirmed this reduction (**Figure 3F**). We next investigated changes in LD accumulation in chondrocytes induced by si-FGFR1 in the presence of FGF8 (**Figure 3G**), and the results revealed that LD accumulation was significantly lower in chondrocytes treated with si-FGFR1 than in those treated with FGF8. Quantification of the number of visible LDs (**Figure 3H**) and total fluorescence intensity (**Figure S11**) further confirmed the changes in LD accumulation in chondrocytes treated with si-FGFR1 in the presence of FGF8.

**Figure 3.**
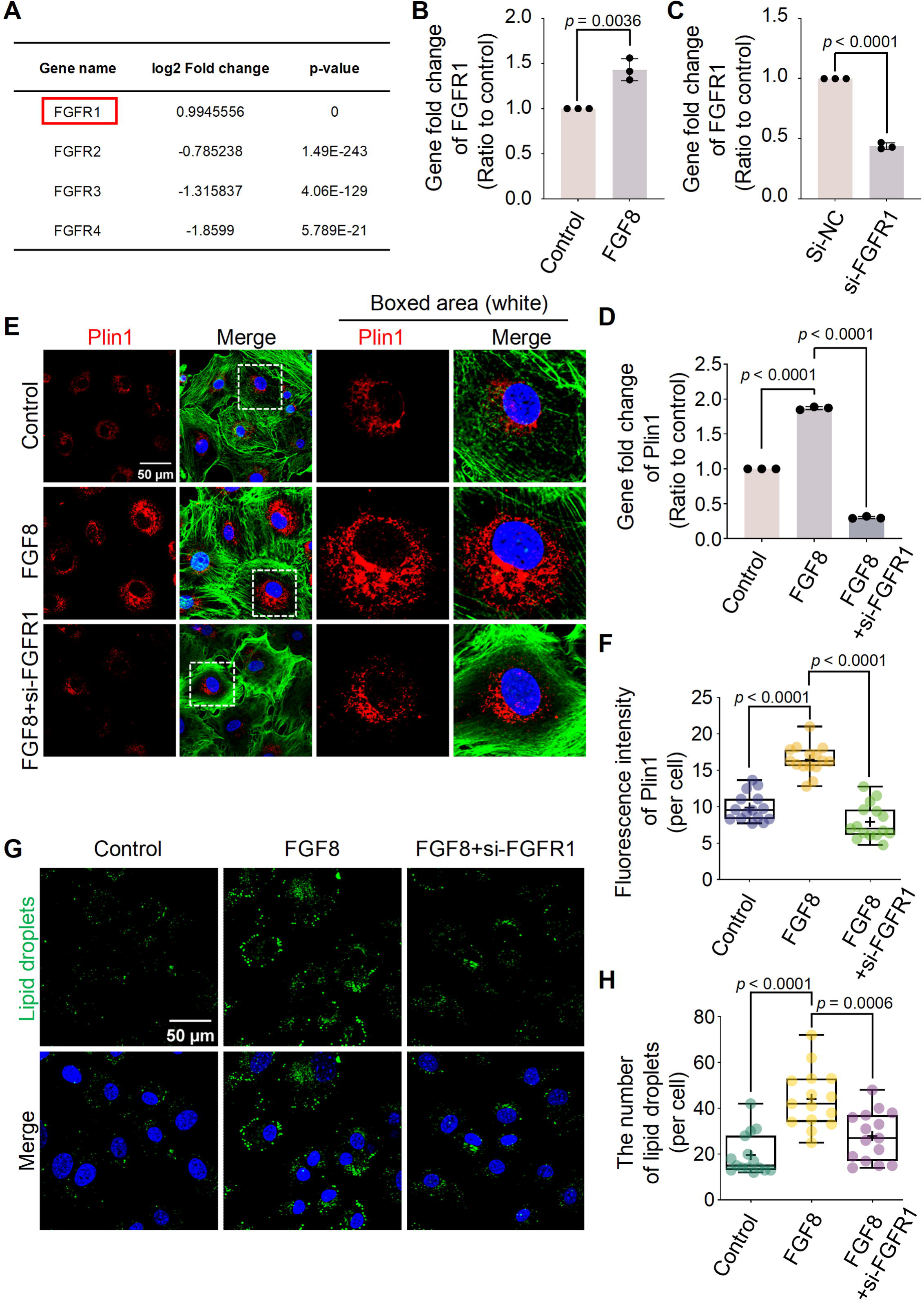
FGF8 enhances lipid droplet accumulation in chondrocytes via FGFR1. (A) RNA sequencing data showing changes in gene expression of FGFRs in chondrocytes exposed to 25 ng/ml FGF8. The red box indicated the upregulated FGFR1 expression. (B) qPCR verifying the expression of FGFR1 in chondrocytes exposed to FGF8 at 25 ng/ml. The data were based on three independent repetitions (n = 3). (C) qPCR verifying the knockdown efficiency of 50 nM si-FGFR1 in chondrocytes. The data were based on three independent experiments (n = 3). (D) qPCR verifying the decreased expression of Plin1 in chondrocytes induced by si-FGFR1 in the presence of FGF8. Chondrocytes were pretreated with siRNA for 12 h and then treated with FGF8 (25 ng/ml) for 12 h. The data were based on three independent experiments (n = 3). (E) Representative fluorescence images showing the expression of Plin1 in chondrocytes exposed to si-FGFR1 in the presence of FGF8. Chondrocytes were pretreated with siRNA for 12 h and then treated with FGF8 at 25 ng/ml for 2 days. The images were captured using CLSM (60 ×). Red fluorescence indicated Plin1, green fluorescence indicated the cytoskeleton (F-actin), and blue fluorescence indicated the nuclei. The white dashed boxes indicated the magnified regions. (F) Total fluorescence quantification per cell, confirming the decrease in Plin1 protein in chondrocytes caused by si-FGFR1 in the presence of FGF8 at 25 ng/ml. The data were based on 15 cells from three independent samples (n = 3). (G) Representative fluorescence images (60 ×) showing the change in lipid droplet accumulation in chondrocytes exposed to si-FGFR1 in the presence of FGF8 at 25 ng/ml. Green indicated the lipid droplets, and blue indicated the nuclei. (H) Quantification of the number of visible lipid droplets per cell confirming the changes in the number of lipid droplets in chondrocytes exposed to si-FGFR1 in the presence of FGF8 at 25 ng/ml. The data were based on 15 cells from three independent replicates (n = 3). The data in **B** and **C** were analyzed using two-tailed Student’s t-tests. The data in **D**, **F**, and **H** were analyzed using one-way analysis of variance. Data in **B**, **C**, and **D** were presented as the means ± SDs. Data in **F** and **H** were shown in the box (25%, 50% to 75%) and whisker (minimum to maximum) plots. Differences were considered statistically significant at p < 0.05.

### 4. FGF8 activates p-p38 signaling in chondrocytes

To further determine which cytoplasmic signaling pathway was activated in chondrocytes exposed to FGF8, we first performed western blotting (**Figure 4A**) and found that FGF8 increased the expression of p-Erk (up to 1.5-fold) and p-p38 (up to 3.0-fold) relative to that in the control groups (**Figure 4B**). Given that the change in p-p38 was greater than that in p-Erk expression in chondrocytes exposed to FGF8, we focused on the role of p-p38 signaling in FGF8-mediated LD accumulation. Immunofluorescence was used to detect the cytoplasmic expression and distribution of p-p38 in chondrocytes exposed to FGF8 (**Figure 4C**), and the results revealed that FGF8 increased the protein expression of p-p38. More importantly, the increase in p-p38 was highly concentrated in the nuclear region of chondrocytes treated with FGF8 at 25 ng/ml (**Figure 4C**, boxed areas). Using total fluorescence quantification per cell (**Figure 4D**) and linear fluorescence quantification in the nuclear region (**Figure 4E**), we further confirmed the increase in p-p38 in chondrocytes exposed to FGF8.

**Figure 4.**
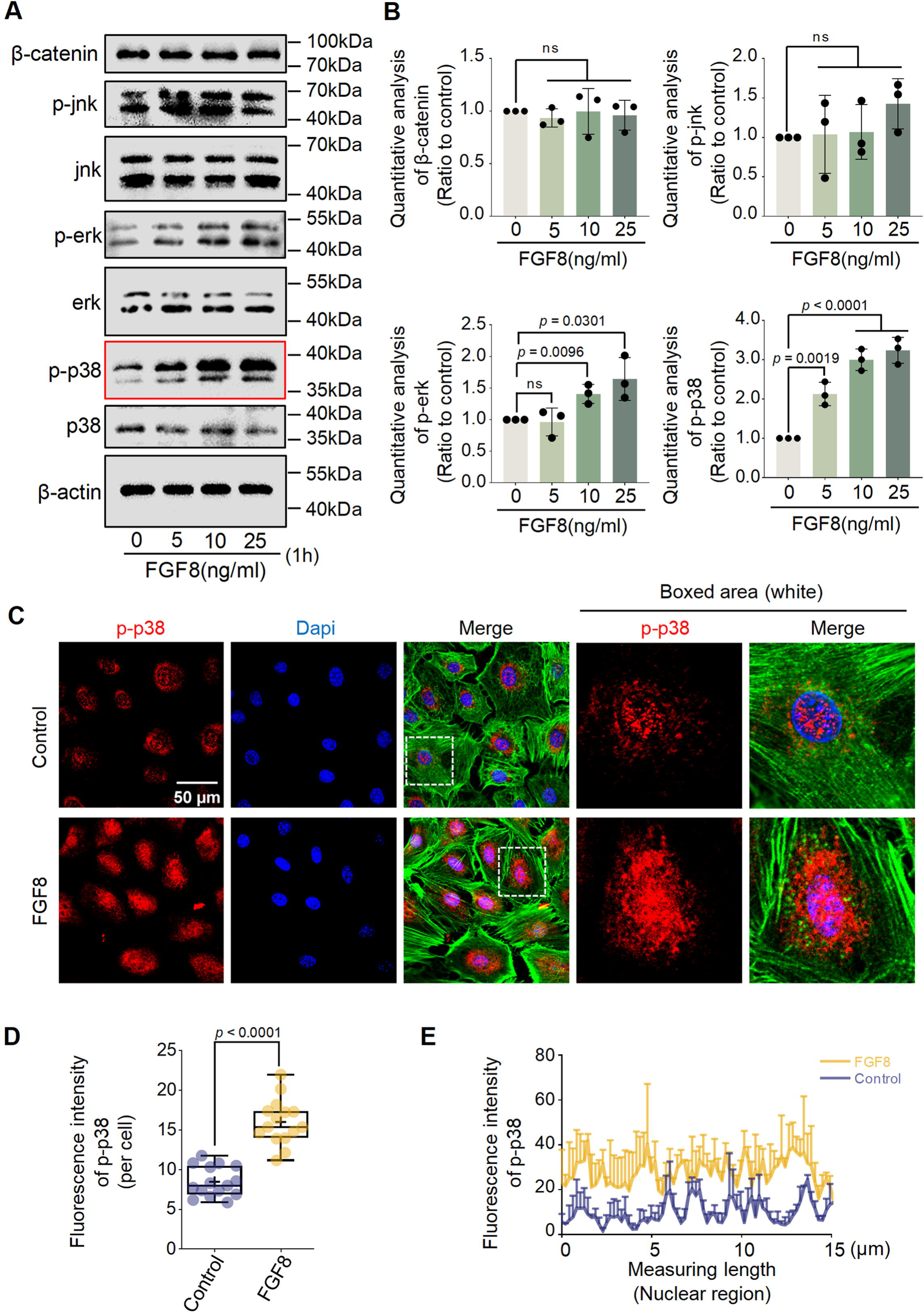
FGF8 activates p-p38 signaling in chondrocytes. **(A)** Western blotting showing upregulation of p-p38 signaling in chondrocytes exposed to different concentrations of FGF8. β-actin was used as the internal reference. The cell lysates were collected after treatment with FGF8 for 1 h. The red box showed the change in p-p38 signaling. Data were representative of three independent samples (n = 3). **(B)** Quantitative analysis confirming the fold-change in p-p38 protein expression in chondrocytes treated with FGF8 in (A). The data were based on three replicates (n = 3). **(C)** Representative fluorescence images (60 ×) showing the change in p-p38 in chondrocytes exposed to FGF8 at 25 ng/ml for 1 h. Red fluorescence indicated p-p38, green fluorescence indicated the cytoskeleton (F-actin), and blue fluorescence indicated the nuclei. The white dashed boxes indicated the magnified regions. **(D)** Total fluorescence quantification per cell confirming the change in p-p38 protein levels in chondrocytes exposed to FGF8 at 25 ng/ml for 1 h. The data were based on 15 cells from three independent samples (n = 3). **(E)** Representative linear fluorescence quantification of chondrocyte nuclei showing a change in p-p38 after exposure to FGF8 at 25 ng/ml. The average nuclear diameter ranged from 8 to 15 µm. The data were based on three independent replicates (n = 3). The data in **B** were based on a one-way analysis of variance. The data in **D** were based on two-tailed Student’s t-tests. Data in **B** were presented as the means ±SDs. Data in **D** were shown in box (from 25%, 50% to 75%) and whisker (minimum to maximum) plots. Differences were considered statistically significant at p < 0.05.

### 5. FGF8 promotes lipid droplet accumulation via the FGFR1/p-p38 axis

To confirm the role of p-p38 signaling in FGF8-mediated LD accumulation, we used SB203580, a specific inhibitor of p-p38 [56]. Western blotting confirmed that SB203580 effectively reduced the upregulation of p-p38 in chondrocytes induced by FGF8 (**Figure 5A & 5B**). Immunofluorescence was used to detect the cytoplasmic expression of p-p38 in chondrocytes exposed to FGF8 (**Figure 5C**), and the results showed that SB203580 treatment significantly reduced the expression of p-p38 in chondrocytes in the presence of FGF8. The total fluorescence quantification per cell (**Figure 5D**) and linear fluorescence quantification in the nuclear region (**Figure 5E**) further confirmed this reduction, thus confirming the effectiveness of blocking p-p38 in the chondrocytes. We then investigated the expression of Plin1 in chondrocytes treated with SB203580 in the presence of FGF8 (**Figure 5F-5H**). qPCR revealed that the gene expression of Plin1 was decreased in chondrocytes with p-p38 blockade in the presence of FGF8 compared to that in the individual FGF8-treated group (**Figure 5F**). Using immunofluorescence, we observed that blocking p-p38 with SB203580 effectively reduced the cytoplasmic expression of Plin1 protein in the presence of FGF8 (**Figure 5G**), and total fluorescence quantification per cell confirmed this reduction (**Figure 5H**). We detected LD accumulation in chondrocytes treated with SB203580 in the presence of FGF8. Using fluorescence, we observed that LD accumulation was significantly reduced in chondrocytes treated with SB203580 in the presence of FGF8, relative to that in the individual FGF8-treated group (**Figure 5I**). Quantification of the number of visible LDs (**Figure 5J**) and total fluorescence intensity (**Figure S12**) further confirmed the changes in LD accumulation in chondrocytes treated with SB203580 in the presence of FGF8.

**Figure 5.**
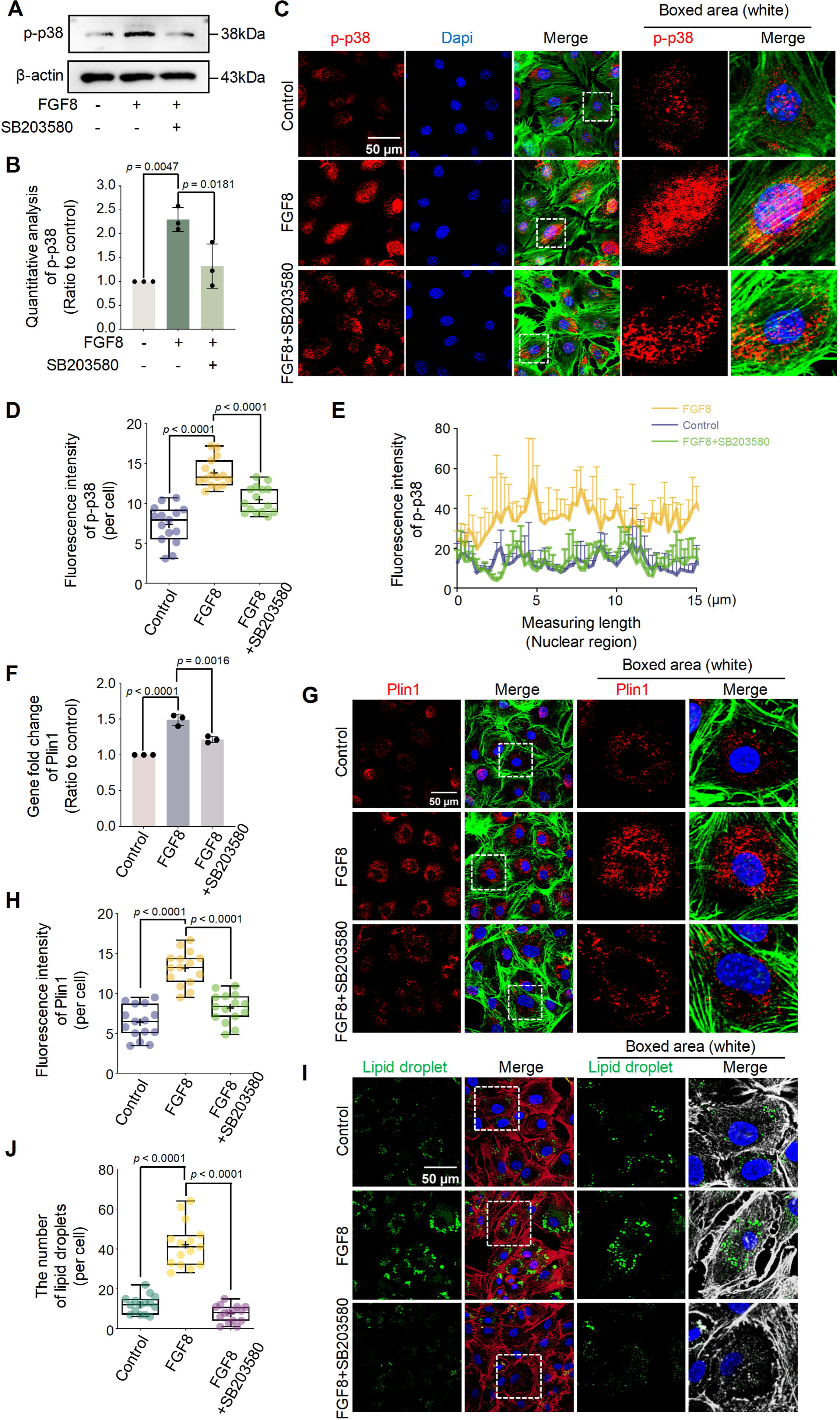
FGF8 promotes lipid droplet accumulation in chondrocytes through p-p38 signaling. **(A)** Western blotting showing that the expression of p-p38 in chondrocytes was inhibited by SB203580 in the presence of FGF8. Chondrocytes were pretreated with SB203580 at 20 µM for 1 h and then treated with FGF8 at 25 ng/ml. β-actin was used as the internal reference. **(B)** Quantitative analysis confirming the change in p-p38 signaling in chondrocytes in (A). The data were based on three replicates (n = 3). **(C)** Representative fluorescence images showing p-p38 protein expression in chondrocytes exposed to SB203580 in the presence of FGF8. Chondrocytes were pretreated with SB203580 at 20 µM for 1 h and then treated with FGF8 at 25 ng/ml. The images were captured using CLSM (60 ×). Red fluorescence indicated Plin1, green fluorescence indicated the cytoskeleton (F-actin), and blue fluorescence indicated the nuclei. The white dashed boxes indicated the magnified regions. **(D)** Total fluorescence quantification per cell showing the change in p-p38 in chondrocytes exposed to SB203580 in the presence of FGF8 at 25 ng/ml. The data were based on 15 cells from three independent samples (n = 3). **(E)** Representative linear fluorescence quantification of chondrocyte nuclei showing the change in nuclear accumulation of p-p38 in chondrocytes inhibited by SB203580 in the presence of 25 ng/ml FGF8. The average nuclear length was in the range of 8 to 15 µm. The data were based on three replicates (n = 3). **(F)** qPCR showing the change in the gene expression of plin1 in chondrocytes exposed to SB203580 in the presence of FGF8. Chondrocytes were pretreated with SB203580 at 20 µM for 1 h and then treated with FGF8 at 25 ng/ml for 12 h. Data were based on three replicates (n = 3). **(G)** Representative fluorescence images showing the change in Plin1 expression in chondrocytes exposed to SB203580 in the presence of FGF8 at 25 ng/ml. The images were obtained using CLSM (60 ×). Red fluorescence indicated Plin1, green fluorescence indicated the cytoskeleton (F-actin), and blue fluorescence indicated the nuclei. The white dashed boxes showed the magnified regions. **(H)** Total fluorescence quantification per cell confirming the change in Plin1 in chondrocytes induced by SB203580 in the presence of FGF8 at 25 ng/ml. The data were based on 15 cells from three independent samples (n = 3). **(I)** Representative fluorescence images showing the change in lipid droplets in chondrocytes induced by SB203580 in the presence of FGF8 at 25 ng/ml. Images were captured using CLSM (60 ×). Green fluorescence indicated lipid droplets, red fluorescence indicated the cytoskeleton (F-actin), and blue fluorescence indicated the nuclei. The white dashed boxes indicated the magnified regions. **(J)** Quantification of the number of visible lipid droplets per cell showing the change in the number of lipid droplets in chondrocytes induced by SB203580 in the presence of 25 ng/ml FGF8. The data were based on 15 cells from three independent samples (n = 3). The data in **B**, **D**, **F**, **H**, and **J** were based on a one-way analysis of variance. The data in **B** and **F** were presented as the means ±SDs. The data in **D**, **H**, and **J** were shown in the box (from 25%, 50% to 75%) and whisker (minimum to maximum) plots. Differences were considered statistically significant at p < 0.05.

To determine the regulatory correlation between FGFR1 and p-p38 in chondrocytes exposed to FGF8, we detected the expression of p-p38 signaling in chondrocytes using siRNA targeting FGFR1 in the presence of FGF8. Using western blotting, we observed that the knockdown of FGFR1 significantly decreased the protein expression of p-p38 in chondrocytes in the presence of FGF8 (**Figure 6A & 6B**). Using immunofluorescence, we found that si-FGFR1 reduced the protein expression and nuclear accumulation of p-p38 in chondrocytes in the presence of FGF8 (**Figure 6C**). Total fluorescence quantification (**Figure 6D**) and linear fluorescence quantification of p-p38 in the nuclear region (**Figure 6E**) further confirmed this result.

**Figure 6.**
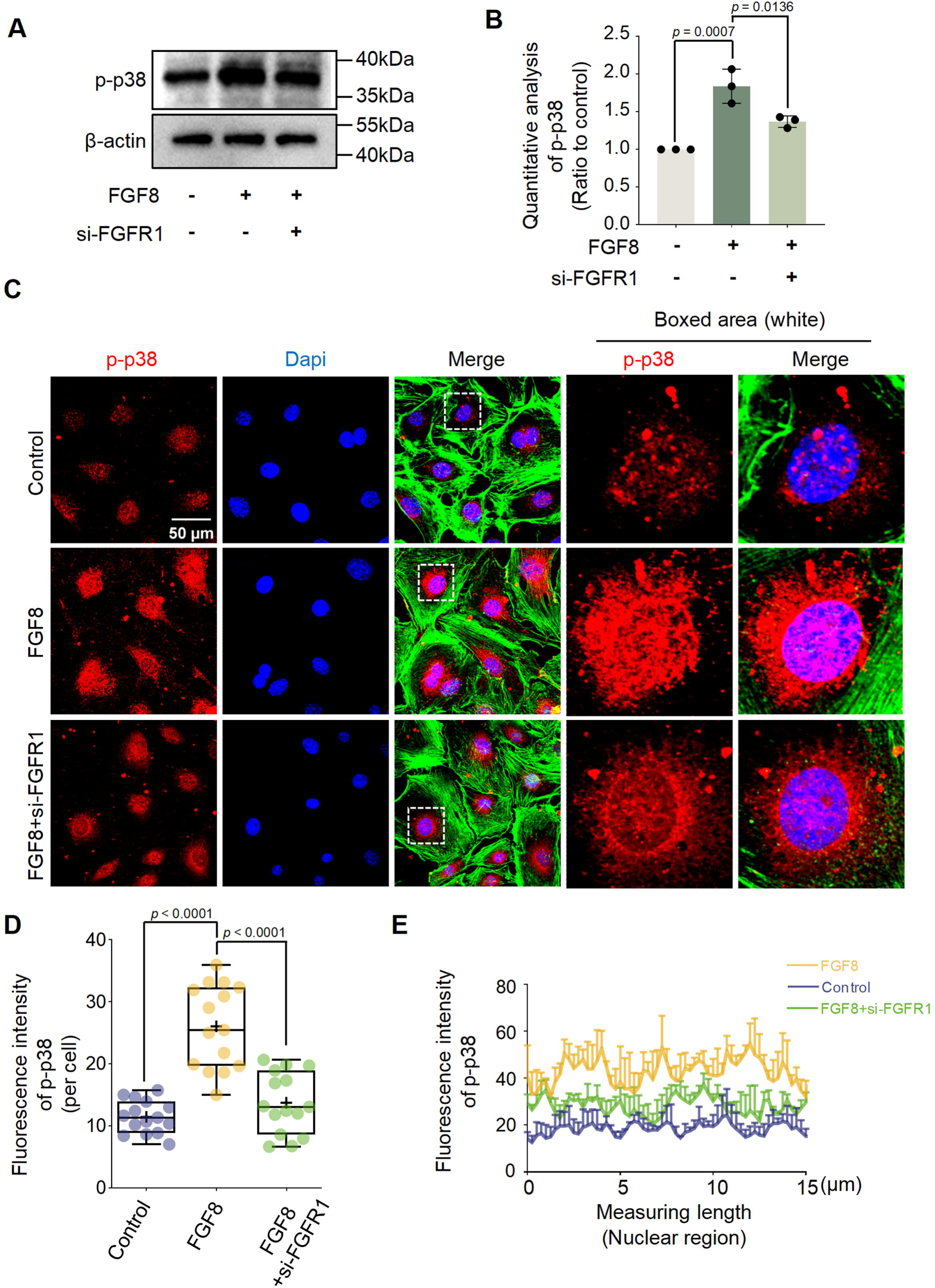
Knockdown of FGFR1 downregulates p-p38 signaling. **(A)** Western blotting showing the change in p-p38 signaling in chondrocytes induced by si-FGFR1 in the presence of FGF8. Chondrocytes were pretreated with siRNA for 12 h and then treated with FGF8 at 25 ng/ml for 1 h. β-actin was used as an internal reference. **(B)** Quantitative analysis confirming the protein change in p-p38 in chondrocytes induced by si-FGFR1 in the presence of FGF8 at 25 ng/ml in (A). The data were based on three independent replicates (n = 3). **(C)** Representative fluorescence images showing the change in p-p38 levels in chondrocytes induced by si-FGFR1 in the presence of FGF8. Chondrocytes were pretreated with siRNA for 12 h and then treated with FGF8 at 25 ng/ml for 1 h. Images were obtained using CLSM (60 ×). Red fluorescence indicated p-p38, green fluorescence indicated the cytoskeleton (F-actin), and blue fluorescence indicated the nuclei. The white dashed boxes indicated the magnified regions. **(D)** Total fluorescence quantification per cell showing the change in p-p38 in chondrocytes induced by si-FGFR1 in the presence of FGF8 at 25 ng/ml. The data were based on 15 cells from three independent samples. **(E)** Representative linear fluorescence quantification of chondrocyte nuclei showing the change in nuclear accumulation of p-p38 induced by si-FGFR1 in the presence of FGF8 at 25 ng/ml. The average nuclear length was in the range of 8 to 15 µm. The data were based on three independent replicates (n = 3). The data in **B** and **D** are based on a one-way analysis of variance. Data in **B** are presented as the means ±SDs. Data in **D** are shown in box (from 25%, 50% to 75%) and whisker (minimum to maximum) plots. Differences were considered statistically significant at p < 0.05.

Collectively, these results indicate that FGF8 promotes LD accumulation in chondrocytes mainly via the FGFR1/p38 axis.

## Discussion

As the only cell type in cartilage, chondrocytes exhibit abundant lipid deposition and LD accumulation [36]. LD accumulation and metabolism are influenced by various biochemical factors [42,43]. FGF8, one of the most important biochemical factors that can modulate the proliferation, differentiation, migration, and metabolism of chondrocytes, has attracted increasing attention in the physiology and pathology of cartilage [44,57]. To date, no study has explored the importance of FGF8 in LD accumulation in chondrocytes. Here, we showed that FGF8 promoted LD accumulation in chondrocytes by upregulating the Plin1 protein and that FGF8-mediated LD accumulation occurred mainly through the FGFR1/p-p38 axis (**Figure 7**). These results expand our understanding of how cellular lipid metabolic homeostasis is remodeled by biochemical factors.

**Figure 7.**
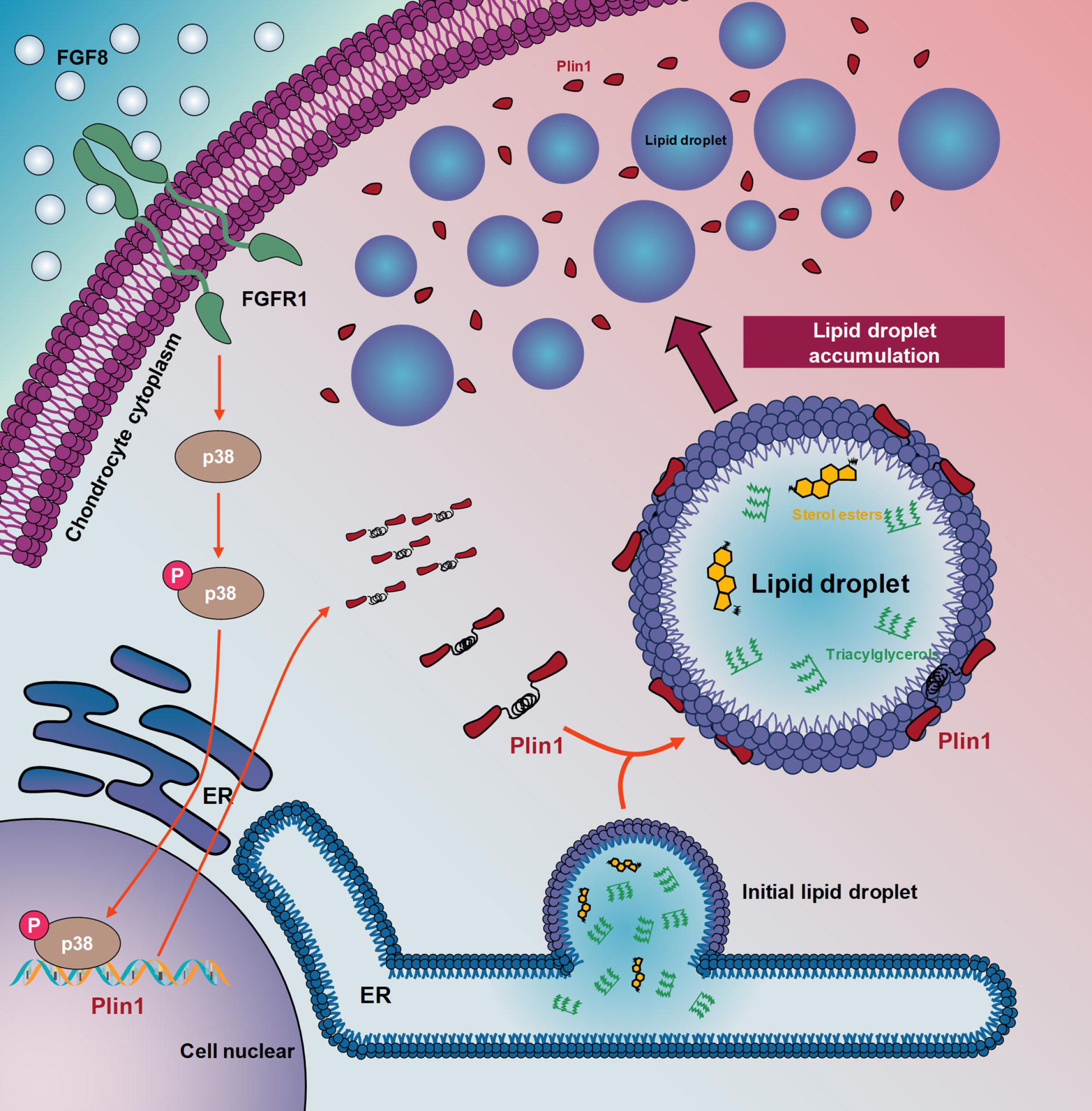
Schematic diagram illustrating the regulatory mechanism of lipid droplet accumulation in chondrocytes following exposure to FGF8. FGF8 binds to FGFR1 and activates p-p38 signaling in chondrocytes. Downstream signaling promotes the expression of plin1, which facilitates the encapsulation of lipid droplets and ultimately increases lipid droplet accumulation in chondrocytes.

Owing to the nutrient-limited microenvironment without the supply of blood vessels, lymphatic vessels, and nerves, cartilage obtains nutrients from the surrounding tissues through diffusion [24,26,58], and its metabolism is thus greatly influenced by neighboring signals [59]. The synovium, a thin layer of connective tissue, provides a nonadherent surface for the joint cartilage and secretes paracrine factors that influence articular chondrocyte metabolism [59,60]. We previously showed that crosstalk with osteoblasts from subchondral bone also increased cartilage lipid catabolism by impairing cholesterol synthesis and accumulation [61] and induced glucose-derived ATP perturbations in chondrocytes [62]. Physiologically, chondrocytes must effectively utilize diffused nutrients, including lipids, to meet energy needs, and thus maintain cellular activities such as signal transmission, protein synthesis, resting potential maintenance, and DNA replication and repair [34]. From a disease perspective, abnormal lipid accumulation in chondrocytes has been associated with a variety of osteoarticular diseases, including OA [37,63]. For example, Park et al. reported that LD accumulation within chondrocytes induced by PPARα and ACOT12 deficiency accelerates the degeneration of the cartilage matrix in OA [30]. Wang et al. reported that GDF11 inhibits abnormal lipid formation and LD accumulation by promoting ubiquitination of PPARγ in chondrocytes in temporomandibular joint osteoarthritis [31]. Meanwhile, as a typical inflammatory disease, OA progression is accompanied by significant upregulation of cytokines in the joint cavity, including FGF8 [44]. Here, we found that FGF8 upregulated multiple lipid metabolism-related pathways, such as fat absorption and digestion, ether lipid metabolism, and arachidonic acid metabolism pathways (**Figure 1A**), and facilitated LD accumulation in chondrocytes (**Figure 1B-1F & S1**). Collectively, the upregulation of lipid accumulation in chondrocytes by FGF8 may explain lipid accumulation and pathological deterioration in osteoarthritic cartilage [47,64,65].

Plin family members, including Plin1-5, are the main modulators of LD accumulation [20]. Compared to other members, Plin1 is a more extensively studied protein that is thought to be responsible for LD accumulation [12]. For example, Wei et al. reported that overexpression of Plin1 blocked LD degradation and promotes abnormal LD growth [66]. Sun et al. showed that Plin1 promotes unilocular lipid droplet formation through the activation of Fsp27 in adipocytes [67]. Wang et al. found that Plin1 formed a complex with apol6, which prevented Plin1 from binding to HSL, thereby inhibiting lipolysis and increasing LD accumulation [68]. Under normal physiological conditions, Plin1 is located on the surface of LDs via its hydrophobic structural domains and binds to α/β-hydrolase domain-containing protein 5 (ABHD5) to inhibit phospholipid lipolysis, thereby promoting LD formation and enlargement [20]. Conversely, during intracellular lipolysis, Plin1 undergoes phosphorylation and facilitates the release of phosphorylated ABHD5. Phosphorylated ABHD5 combines with phosphorylated adipose triglyceride lipase (ATGL) to further enhance lipolysis [20]. In addition, phosphorylated Plin1 recruits and binds to HSL to increase lipolysis [20]. For example, Cao et al. reported that with a high-fat diet, Plin1 could be phosphorylated by adenylate cyclase 7 and promote inflammatory lipolysis of LDs in fibroblast-like synoviocytes, exacerbating OA [64]. By knocking down and overexpressing of Plin1, we found that LD accumulation also changed significantly with the changes in Plin1 expression (**Figure 2**). These results confirmed that Plin1 plays an important role in the accumulation of LDs in chondrocytes. Furthermore, we also detected the expression of HSL, DGAT2, and ATGL in chondrocytes induced by FGF8, and found that the expression of DGAT2 and ATGL was not affected but that of HSL was significantly reduced (**Figure S4**). HSL is a vital lipolytic enzyme that hydrolyzes triglycerides [70]. There is a reason to infer that their combined effect on the increase in Plin1 and decrease in HSL enhances lipid accumulation and LD formation. Notably, Plin1 is not the only factor that regulates lipid accumulation in chondrocytes, and many other factors are involved, as shown in **Figure 2A & S3**. For example, apolipoprotein L6 (apol6) was upregulated in chondrocytes exposed to FGF8 [68]. Previous reports have shown that apol6 can form a complex with Plin1, which prevents Plin1 from binding to HSL, thereby inhibiting lipolysis and increasing LD accumulation in the adipose tissue. Taken together, these results indicate that a series of regulatory protein mediators, including Plin1, promotes lipid accumulation and LD formation induced by FGF8.

FGF signals enter the cellular cytoplasm mainly via FGF receptors (FGFR1-4) [57,71]. Chondrocytes in different zones of cartilage spatiotemporally express FGFRs [72]. During cartilage development, chondrocytes highly express FGFR3 in the superficial zone, and FGFR1 and 2 in the hypertrophic and calcified zones [64]. These receptors selectively recognize their FGF ligands, thereby allowing the entry of FGF signals to regulate the physiological and pathological behaviors of chondrocytes. For example, FGF2 increases the catabolic activity of human articular chondrocytes via FGFR1 [73–75]. FGF18 has been shown to facilitate chondrogenesis through FGFR3 [76]. Yellapragada et al. reported that FGF8 binds to FGFR1 to regulate the differentiation of hypothalamic neurons that express gonadotropin-releasing hormones [77]. Jacques et al. demonstrated that FGF8 binds to FGFR3 to promote cochlear development in mammals [78]. In chondrocytes, we found that FGF8 mediates LD accumulation via FGFR1 (**Figure 3 & S10**). Given that FGFR1 expression is elevated in cartilage during OA progression, and its inhibition attenuates OA-associated inflammation [48,64], the potential role of FGFR1 in OA pathogenesis warrants attention because of its ability to target LD accumulation.

Previous studies have reported that FGFs relay signals to cytoplasmic MAPK, PI3K/AKT, NF-κB, or β-catenin/Wnt signaling to regulate target gene expression [44,64,69]. For example, FGF2 has been shown to mediate the elongation of primary cilia in chondrocytes mainly through Erk signaling [79], and to modulate chondrocyte differentiation through crosstalk between Erk and β-catenin/WNT signaling [80]. FGF23 has been reported to accelerate chondrocyte hypertrophy and degeneration of the OA cartilage matrix via β-catenin/WNT signaling [81]. Lin et al. reported that FGF8 activates PI3K/AKT and p-p38 signaling to regulate the pace of tooth development [82]. FGF8 can also activate β-catenin/WNT signaling to induce the differentiation of dental mesenchymal cells into odontoblast-like cells [83]. We previously reported that FGF8 activates NF-κB signaling to upregulate the expression of matrix metalloproteinases 2 and 9 (MMP-2 & -9) in chondrocytes [48]. In this study, we showed that FGF8 activates MAPKs, including Erk and p38 in chondrocytes (the expression of p38 was much greater than that of Erk, **Figure 4A & 4B**) and revealed the importance of p-p38 signaling in FGF8-mediated LD accumulation via its inhibition with SB203580 (**Figure 5**). These findings are consistent with reports that MAPK/p38 signaling is regulated by FGF8 [56]. In addition, some reports have correlated p-p38 signaling with lipid metabolism. Liu et al. demonstrated that the upregulation of p-p38 signaling in macrophages promoted lipid accumulation following activation by reactive oxygen species [84]. Li et al. reported that upregulation of p-p38 signaling in HCT-116 cancer cells enhanced lipid accumulation by promoting triacylglycerol biosynthesis [85]. In this study, we established an association between p-p38 and LD accumulation via Plin1 protein in chondrocytes.

In conclusion, in the present study, we found that FGF8 promoted LD accumulation in chondrocytes by increasing Plin1 expression through FGFR1/p38 signaling. These findings improve our understanding of LDs in chondrocytes and provide potential therapeutic targets for cartilage diseases.

## Supplementary Data

Supplementary data are presented in Supplementary Information file.

## Supporting information

Supplementary materials for original paper

## Acknowledgments

None

## Data Availability Statement

All data generated in this study are available from the corresponding author upon request.

