## Supplementary materials for original paper for "FGF8 promotes lipid droplet accumulation via the FGFR1/p-p38 axis in chondrocytes"

**FGF8 dominates lipid droplet accumulation via FGFR1/p-p38 signaling in chondrocytes**

### 1. Supplementary figures

**Figure S1**

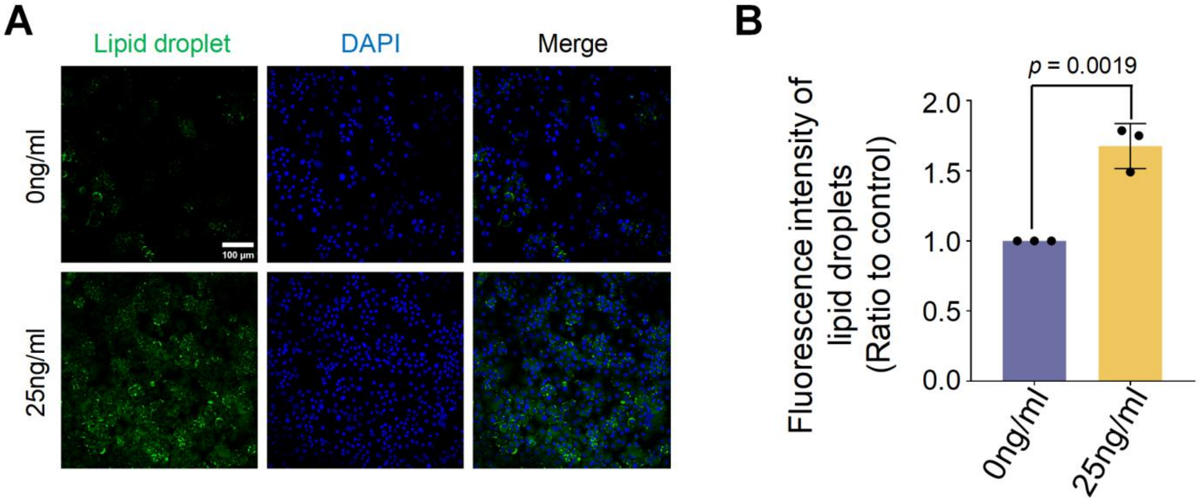


**Figure S1. FGF8 promotes lipid droplet accumulation in chondrocytes.**

**(A)** Representative fluorescence images (40×) showing lipid droplet accumulation in chondrocytes induced by FGF8 at 25 ng/ml for 2 days.

**(B)** Total fluorescence quantification showing the change of lipid droplets in chondrocytes induced by FGF8 at 25 ng/ml for 2 days in (A). The data are based on 3 independent replicates (n = 3).

The data in B are based on two-tailed Student's t test. The data in B are presented as mean ± SD. Differences are considered significant at $p < 0.05$.

**Figure S2**

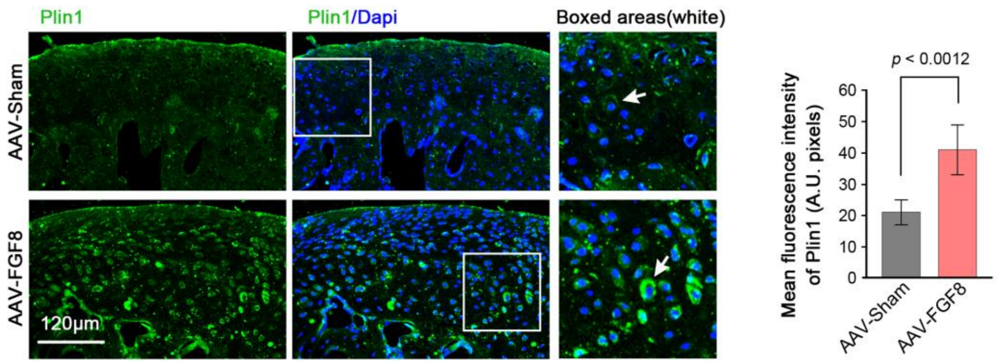


**Figure S2. FGF8 induced a higher expression of Plin1 in cartilage tissue.** The white arrows indicate the representative expressions of Plin1 in chondrocytes. This figure was related to the figure 2C.

**Figure S3**

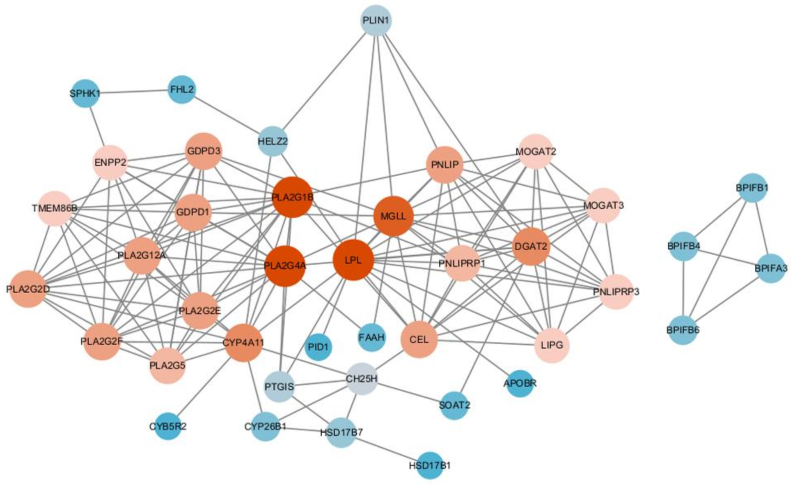


**Figure S3. A protein-protein interaction analysis using STRING confirmed that FGF8-mediated lipid accumulation involved a series of regulatory proteins.** This figure was related to Figure 2A.

**Figure S4**

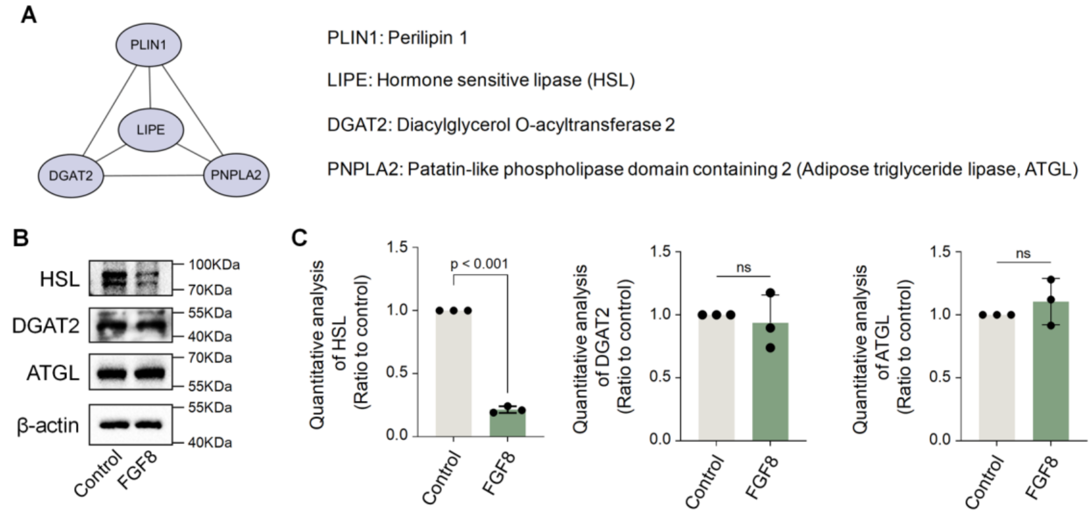


**Figure S4. The expressions of Plin1 partner proteins in chondrocytes induced by FGF8.** (A) A protein-protein interaction analysis by STRING confirmed the relationship between Plin1 and the partner proteins. (B) Western blotting showed changes in HSL, DGAT2, and ATGL in chondrocytes induced by FGF8. (C) Quantitative analysis of HSL, DGAT2, and ATGL in (b). The data are based on 3 independent results (n = 3).

**Figure S5**

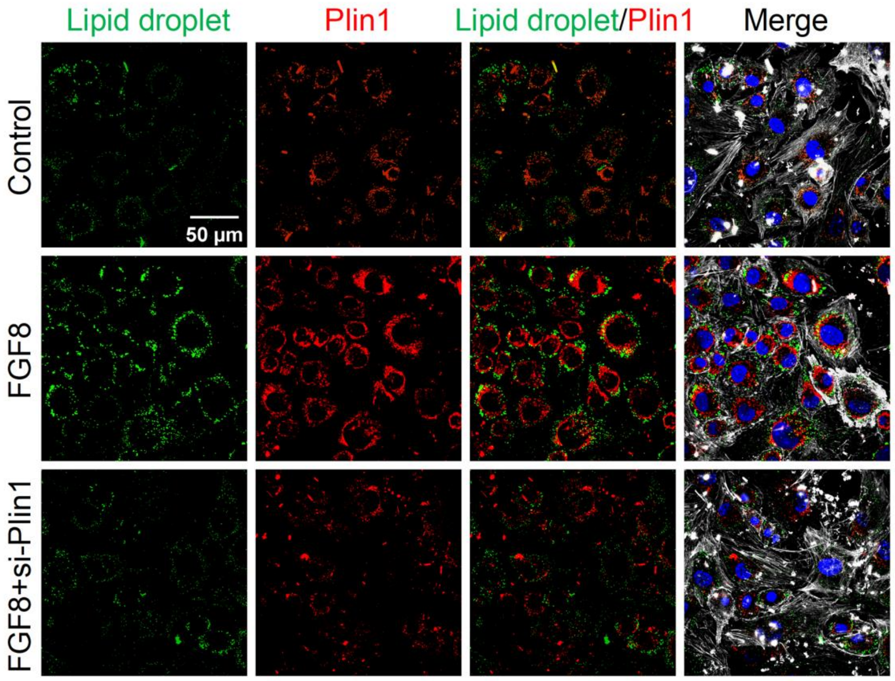


**Figure S5. FGF8 promotes lipid droplet accumulation and plin1 expression in chondrocytes.**

Representative fluorescence images (60×) of multiple cells showing the changes of lipid droplets and plin1 in chondrocytes induced by si-plin1 in the presence of FGF8.

**Figure S6**

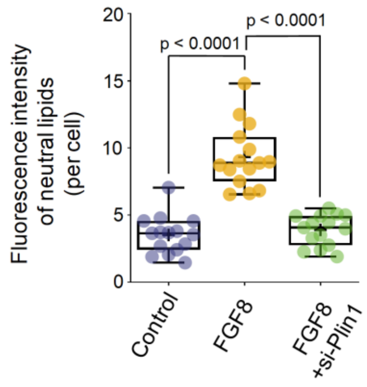


**Figure S6. Total fluorescence quantification of lipids per cell by si-Plin1 in the presence of FGF8.**

The data are based on 15 cells from 3 independent samples (n = 3). This figure is related to figure 2F. The data are shown in the box (from 25%, 50% to 75%) and whisker (minimum to maximum) plots. The statistic analysis is based on one-way analysis of variance and differences are considered significant at $p < 0.05$.

**Figure S7**

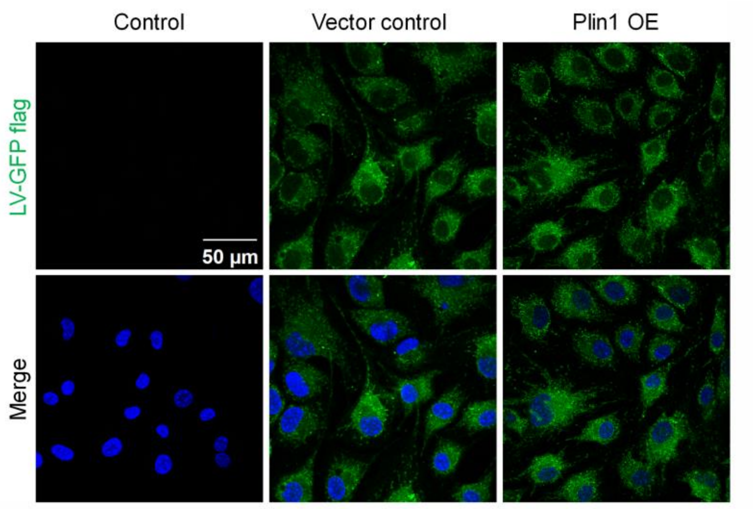


**Figure S7. The transfection efficiency of lentivirus carrying Plin1 gene in chondrocytes.**

Representative fluorescence images (60×) of multiple cells showing the the efficiency of lentivirus transfection in chondrocytes at 30 MOI (multiplicity of infection). Green fluorescence indicated GFP protein carried by lentivirus and blue fluorescence indicated the nuclei.

**Figure S8**

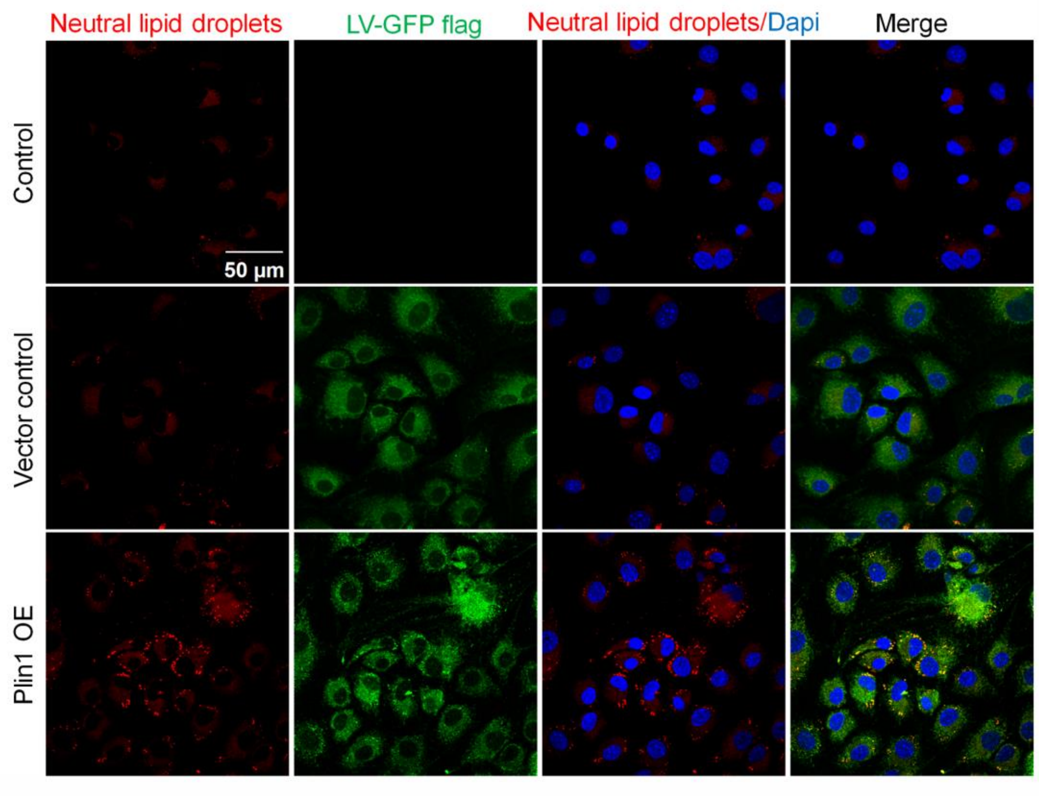


**Figure S8. Overexpression of Plin1 promotes lipid droplet accumulation in chondrocytes.**

Representative fluorescence images (60×) of multiple cells showing the changes of lipid droplet accumulation in chondrocytes induced by overexpressing Plin1.

**Figure S9**

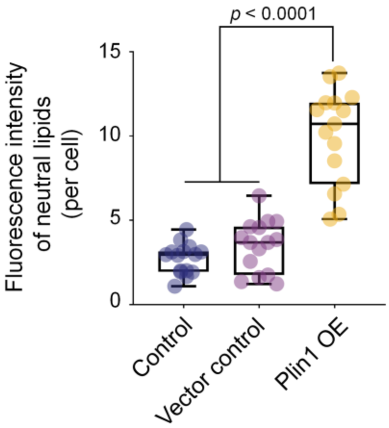


**Figure S9. Total fluorescence quantification of lipids per cell by overexpression of Plin1.**

The data are based on 15 cells from 3 independent samples (n = 3). This figure is related to figure 2P. The data are shown in the box (from 25%, 50% to 75%) and whisker (minimum to maximum) plots. The statistic analysis is based on two-tailed Student’s t test and differences are considered significant at $p < 0.05$.

**Figure S10**

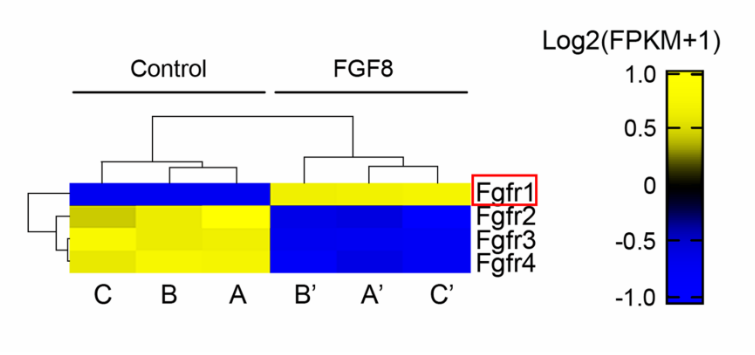


**Figure S10. The gene expression of FGFRs in chondrocytes induced by FGF8.**

The pheatmap based on RNA sequencing showing the gene changes of FGFRs in chondrocytes induced by FGF8 at 25 ng/ml for 24 h.

**Figure S11**

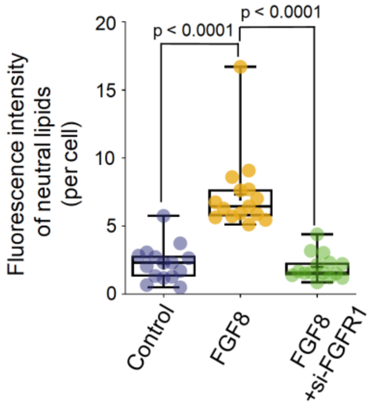


**Figure S11. Total fluorescence quantification of lipids per cell by si-FGFR1 in the presence of FGF8.**

The data are based on 15 cells from 3 independent samples (n = 3). This figure is related to figure 3G. The data are shown in the box (from 25%, 50% to 75%) and whisker (minimum to maximum) plots. The statistic analysis is based on one-way analysis of variance and differences are considered significant at $p < 0.05$.

**Figure S12**

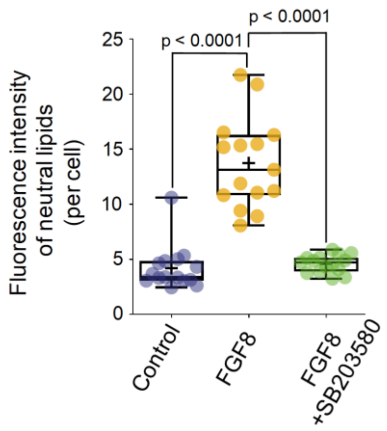


**Figure S12. Total fluorescence quantification of lipids per cell treated with SB203580 in the presence of FGF8.** The data are based on 15 cells from 3 independent samples (n = 3). This figure is related to figure 5I. The data are shown in the box (from 25%, 50% to 75%) and whisker (minimum to maximum) plots. The statistic analysis is based on one-way analysis of variance and differences are considered significant at $p < 0.05$.

### 2. Supplementary tables

**Table S1.** (a) KEGG analysis showed up-regulated pathways in chondrocytes induced by FGF8.

| #Term | Database | ID | Input number | Background number | P-Value | Corrected P-Value |
|---|---|---|---|---|---|---|
| Cytokine-cytokine receptor interaction | KEGG PATHWAY | mmu04060 | 23 | 264 | 6.24218E-10 | 1.35455E-07 |
| TNF signaling pathway | KEGG PATHWAY | mmu04668 | 11 | 110 | 5.84458E-06 | 0.000634137 |
| PI3K-Akt signaling pathway | KEGG PATHWAY | mmu04151 | 16 | 341 | 0.000351566 | 0.010819147 |
| NF-kappa B signaling pathway | KEGG PATHWAY | mmu04064 | 8 | 99 | 0.000424719 | 0.010819147 |
| Renin-angiotensin system | KEGG PATHWAY | mmu04614 | 5 | 34 | 0.00044872 | 0.010819147 |
| Chemokine signaling pathway | KEGG PATHWAY | mmu04062 | 11 | 194 | 0.000679331 | 0.014741489 |
| VEGF signaling pathway | KEGG PATHWAY | mmu04370 | 6 | 60 | 0.000798775 | 0.015757657 |
| Ras signaling pathway | KEGG PATHWAY | mmu04014 | 11 | 228 | 0.00232283 | 0.03877339 |
| EGFR tyrosine kinase inhibitor resistance | KEGG PATHWAY | mmu01521 | 6 | 81 | 0.003318603 | 0.047419579 |
| ECM-receptor interaction | KEGG PATHWAY | mmu04512 | 6 | 83 | 0.003714898 | 0.047419579 |
| Fc gamma R-mediated phagocytosis | KEGG PATHWAY | mmu04666 | 6 | 86 | 0.004373749 | 0.052727976 |
| Thyroid hormone signaling pathway | KEGG PATHWAY | mmu04919 | 7 | 117 | 0.004765717 | 0.052855192 |
| MAPK signaling pathway | KEGG PATHWAY | mmu04010 | 11 | 254 | 0.005066911 | 0.052855192 |
| Arachidonic acid metabolism | KEGG PATHWAY | mmu00590 | 6 | 89 | 0.005115019 | 0.052855192 |
| Ether lipid metabolism | KEGG PATHWAY | mmu00565 | 4 | 44 | 0.008240943 | 0.071802199 |
| Focal adhesion | KEGG PATHWAY | mmu04510 | 9 | 202 | 0.00908307 | 0.073006208 |
| Pancreatic secretion | KEGG PATHWAY | mmu04972 | 6 | 102 | 0.009420156 | 0.073006208 |
| Mucin type O-Glycan biosynthesis | KEGG PATHWAY | mmu00512 | 3 | 29 | 0.015847431 | 0.101143896 |
| Jak-STAT signaling pathway | KEGG PATHWAY | mmu04630 | 7 | 161 | 0.022655585 | 0.140464624 |
| Metabolism of xenobiotics by cytochrome P450 | KEGG PATHWAY | mmu00980 | 4 | 66 | 0.029270346 | 0.171666622 |
| Glycosaminoglycan biosynthesis - keratan sulfate | KEGG PATHWAY | mmu00533 | 2 | 15 | 0.031822453 | 0.17368437 |
| Glycosphingolipid biosynthesis - globo series | KEGG PATHWAY | mmu00603 | 2 | 15 | 0.031822453 | 0.17368437 |
| RIG-I-like receptor signaling pathway | KEGG PATHWAY | mmu04622 | 4 | 68 | 0.032015552 | 0.17368437 |
| Aldosterone-regulated sodium reabsorption | KEGG PATHWAY | mmu04960 | 3 | 40 | 0.034516091 | 0.174185857 |
| Fat digestion and absorption | KEGG PATHWAY | mmu04975 | 3 | 40 | 0.034516091 | 0.174185857 |
| Metabolic pathways | KEGG PATHWAY | mmu01100 | 30 | 1268 | 0.043810445 | 0.211263702 |
| PPAR signaling pathway | KEGG PATHWAY | mmu03320 | 4 | 85 | 0.045262275 | 0.250828558 |
| Regulation of lipolysis in adipocytes | KEGG PATHWAY | mmu04923 | 3 | 57 | 0.048124207 | 0.282549216 |

(b) KEGG analysis showed down-regulated pathways in chondrocytes induced by FGF8.

| #Term | Database | ID | Input number | Background number | P-Value | Corrected P-Value |
|---|---|---|---|---|---|---|
| PI3K-Akt signaling pathway | KEGG PATHWAY | mmu04151 | 25 | 341 | 9.37688E-11 | 1.83787E-08 |
| ECM-receptor interaction | KEGG PATHWAY | mmu04512 | 12 | 83 | 9.46463E-09 | 9.27534E-07 |
| Focal adhesion | KEGG PATHWAY | mmu04510 | 14 | 202 | 2.52556E-06 | 0.000123752 |
| Regulation of actin cytoskeleton | KEGG PATHWAY | mmu04810 | 13 | 216 | 2.37679E-05 | 0.000931703 |
| Rap1 signaling pathway | KEGG PATHWAY | mmu04015 | 12 | 216 | 0.000100568 | 0.003285226 |
| Aldosterone synthesis and secretion | KEGG PATHWAY | mmu04925 | 7 | 85 | 0.000321622 | 0.009005411 |
| MAPK signaling pathway | KEGG PATHWAY | mmu04010 | 12 | 254 | 0.000415908 | 0.010189751 |
| Longevity regulating pathway | KEGG PATHWAY | mmu04211 | 7 | 95 | 0.000601707 | 0.013103836 |
| Glucagon signaling pathway | KEGG PATHWAY | mmu04922 | 7 | 100 | 0.000800494 | 0.015689689 |
| FoxO signaling pathway | KEGG PATHWAY | mmu04068 | 8 | 134 | 0.000923594 | 0.016456763 |
| Glycine, serine and threonine metabolism | KEGG PATHWAY | mmu00260 | 4 | 40 | 0.003345889 | 0.043719613 |
| cGMP-PKG signaling pathway | KEGG PATHWAY | mmu04022 | 8 | 172 | 0.004072213 | 0.049884611 |
| Glycosaminoglycan biosynthesis - heparan sulfate / I | KEGG PATHWAY | mmu00534 | 3 | 24 | 0.006274038 | 0.064738376 |
| Ras signaling pathway | KEGG PATHWAY | mmu04014 | 9 | 228 | 0.006605957 | 0.064738376 |
| AMPK signaling pathway | KEGG PATHWAY | mmu04152 | 6 | 127 | 0.011426417 | 0.106646562 |
| Insulin signaling pathway | KEGG PATHWAY | mmu04910 | 6 | 140 | 0.017396111 | 0.133389896 |
| AGE-RAGE signaling pathway in diabetic complicati | KEGG PATHWAY | mmu04933 | 5 | 102 | 0.017694578 | 0.133389896 |
| Non-alcoholic fatty liver disease (NAFLD) | KEGG PATHWAY | mmu04932 | 6 | 147 | 0.021365781 | 0.148465923 |
| Adipocytokine signaling pathway | KEGG PATHWAY | mmu04920 | 4 | 72 | 0.022319216 | 0.148465923 |
| Cholinergic synapse | KEGG PATHWAY | mmu04725 | 5 | 111 | 0.024139254 | 0.148465923 |
| cAMP signaling pathway | KEGG PATHWAY | mmu04024 | 7 | 197 | 0.025755144 | 0.148465923 |
| Hippo signaling pathway | KEGG PATHWAY | mmu04390 | 6 | 154 | 0.025907206 | 0.148465923 |
| mTOR signaling pathway | KEGG PATHWAY | mmu04150 | 6 | 155 | 0.026604387 | 0.148465923 |
| Biosynthesis of amino acids | KEGG PATHWAY | mmu01230 | 4 | 78 | 0.028522047 | 0.148465923 |
| Platelet activation | KEGG PATHWAY | mmu04611 | 5 | 121 | 0.032881002 | 0.16524811 |
| TGF-beta signaling pathway | KEGG PATHWAY | mmu04350 | 4 | 84 | 0.03565273 | 0.166379406 |
| Fc gamma R-mediated phagocytosis | KEGG PATHWAY | mmu04666 | 4 | 86 | 0.038238296 | 0.173554534 |
| Sphingolipid metabolism | KEGG PATHWAY | mmu00600 | 2 | 48 | 0.045597001 | 0.408421074 |
| Steroid biosynthesis | KEGG PATHWAY | mmu00100 | 1 | 19 | 0.046689692 | 0.53394446 |

**Table S2.** Original data for candidate mediators involved in lipid droplet accumulation.

| ensembl_id | gene_name | gene_biotype | log2FoldChange | pvalue | padj | Control1_FPKM | Control2_FPKM | Control3_FPKM | treated1_FPKM | treated2_FPKM | treated3_FPKM |
|---|---|---|---|---|---|---|---|---|---|---|---|
| ENSMUSG00000027485 | Bpifb1 | protein_coding | 6.233028229 | 6.96E-167 | 9.84E-165 | 0.196377891 | 0.258486758 | 0.336002373 | 22.46846158 | 19.83912441 | 20.38498786 |
| ENSMUSG00000074665 | Bpifb4 | protein_coding | 5.256933342 | 3.868E-54 | 1.655E-52 | 0.139784769 | 0.099663853 | 0.108823078 | 4.887479804 | 4.371030379 | 4.672597131 |
| ENSMUSG00000052974 | Cyp2f2 | protein_coding | 2.92518632 | 1.255E-80 | 7.783E-79 | 1.133240299 | 1.077304727 | 0.984259634 | 8.511297014 | 8.298853671 | 8.417981777 |
| ENSMUSG00000050370 | Ch25h | protein_coding | 2.573230631 | 0.0012999 | 0.0059117 | 0.323782943 | 0.046170203 | 0.072018998 | 1.115558346 | 0.539425489 | 1.102354757 |
| ENSMUSG00000056656 | Apol8 | protein_coding | 2.34669003 | 3.159E-13 | 4.137E-12 | 0.184464455 | 0.252517651 | 0.213358104 | 1.073376532 | 0.983422565 | 1.355021423 |
| ENSMUSG00000041193 | Pla2g5 | protein_coding | 2.132472717 | 2.56E-09 | 2.486E-08 | 1.345119411 | 0.540306636 | 0.9776504 | 4.351605894 | 4.350855323 | 4.456468211 |
| ENSMUSG00000041481 | Serpina3g | protein_coding | 2.05350367 | 1.647E-06 | 1.194E-05 | 0.363390595 | 0.148051719 | 0.277127734 | 0.821432512 | 1.277353249 | 1.349676413 |
| ENSMUSG00000061878 | Sphk1 | protein_coding | 1.75733949 | 2.209E-85 | 1.427E-83 | 1.957092907 | 2.258703564 | 2.06711419 | 7.337354855 | 7.09896181 | 7.632558736 |
| ENSMUSG00000040612 | Ildr2 | protein_coding | 1.752748291 | 3.664E-92 | 2.517E-90 | 1.523066957 | 1.442853804 | 1.573097059 | 5.570355578 | 5.322692235 | 5.110298077 |
| ENSMUSG00000052396 | Mogat2 | protein_coding | 1.724265435 | 1.795E-05 | 0.000113 | 0.267981114 | 0.271737432 | 0.476856033 | 1.003090611 | 1.007394907 | 1.523690746 |
| ENSMUSG00000033576 | Apol6 | protein_coding | 1.499346453 | 2.168E-09 | 2.119E-08 | 0.291534219 | 0.318716049 | 0.507959693 | 1.091253162 | 1.120843398 | 1.122896635 |
| ENSMUSG00000030546 | Plin1 | protein_coding | 1.497289446 | 8.084E-09 | 7.497E-08 | 1.539605652 | 0.85805668 | 1.244337703 | 3.683334183 | 3.241265552 | 3.831587594 |
| ENSMUSG00000057346 | Apol9a | protein_coding | 1.405302798 | 6.605E-27 | 1.522E-25 | 1.725518874 | 1.465843825 | 1.380071753 | 4.366287497 | 4.347383701 | 3.916479607 |
| ENSMUSG00000058914 | C1qtnf3 | protein_coding | 1.363117091 | 2.771E-25 | 6.047E-24 | 2.649651183 | 2.26698042 | 2.528551235 | 7.307627025 | 6.353150591 | 6.384384749 |
| ENSMUSG00000039529 | Atp8b1 | protein_coding | 1.361208171 | 2.705E-39 | 8.715E-38 | 1.501706796 | 1.248531177 | 1.411583536 | 3.86288061 | 3.583713053 | 3.73935415 |
| ENSMUSG00000022425 | Enpp2 | protein_coding | 1.338368723 | 2.595E-98 | 1.976E-96 | 13.3972266 | 12.02590369 | 12.68526348 | 34.34120606 | 33.62586656 | 32.60295392 |
| ENSMUSG00000026399 | Cd55 | protein_coding | 1.267508867 | 2.443E-94 | 1.773E-92 | 7.475763182 | 7.462105584 | 7.602953417 | 18.07926313 | 18.66472555 | 19.8208271 |
| ENSMUSG00000053846 | Lipg | protein_coding | 1.229635833 | 1.096E-08 | 1.005E-07 | 1.183085944 | 1.810476653 | 1.553248383 | 4.035687707 | 3.180277631 | 3.870163073 |
| ENSMUSG00000056220 | Pla2g4a | protein_coding | 1.096428372 | 5.701E-54 | 2.434E-52 | 3.017891592 | 3.062360957 | 2.74592853 | 6.467131393 | 6.556237238 | 6.597569932 |
| ENSMUSG00000029822 | Osbpl3 | protein_coding | 1.049187402 | 1.177E-40 | 3.924E-39 | 2.18650395 | 2.224829516 | 2.247150797 | 4.581691247 | 5.098123526 | 4.699096674 |
| ENSMUSG00000013846 | St3gal1 | protein_coding | 1.045978948 | 3.08E-168 | 4.42E-166 | 17.69705831 | 17.86829297 | 17.58321067 | 36.91389244 | 38.21700247 | 39.23137242 |
| ENSMUSG00000027580 | Helz2 | protein_coding | 1.001553226 | 1.401E-65 | 7.245E-64 | 4.279508155 | 3.82769361 | 4.052122756 | 8.442403134 | 8.44461808 | 8.527192425 |
| ENSMUSG00000055725 | Paqr3 | protein_coding | -1.102090307 | 3.712E-13 | 4.834E-12 | 2.094254263 | 1.631151884 | 1.869723975 | 0.943639098 | 0.925476842 | 0.858296545 |
| ENSMUSG00000045658 | Pid1 | protein_coding | -1.197554947 | 4.25E-64 | 2.157E-62 | 6.204651103 | 6.147552305 | 6.073628567 | 2.859880979 | 2.570542505 | 2.9302682 |
| ENSMUSG00000023045 | Soat2 | protein_coding | -1.45801153 | 5.442E-11 | 6.093E-10 | 3.407182682 | 3.323503512 | 3.734378659 | 1.537484391 | 1.493456157 | 0.97814003 |
| ENSMUSG00000017969 | Ptgis | protein_coding | -1.459310324 | 1.025E-90 | 6.931E-89 | 37.24859768 | 39.57831352 | 39.08720975 | 13.53706369 | 14.72989951 | 15.6177009 |
| ENSMUSG00000048065 | Cyb5r2 | protein_coding | -1.68155176 | 1.01E-50 | 4.078E-49 | 8.160276273 | 7.847435407 | 8.540564566 | 2.937797954 | 2.384573214 | 2.668036896 |
| ENSMUSG00000019301 | Hsd17b1 | protein_coding | -1.748060611 | 2.195E-14 | 3.082E-13 | 4.406978439 | 4.096355539 | 3.920974551 | 1.291348347 | 1.631570746 | 0.976694248 |
| ENSMUSG00000042759 | Apobr | protein_coding | -1.859077593 | 5.211E-57 | 2.379E-55 | 6.144828768 | 6.020208342 | 6.419105281 | 1.653116364 | 1.64538664 | 2.016745152 |
| ENSMUSG00000008136 | Fhl2 | protein_coding | -1.885912345 | 8.555E-57 | 3.865E-55 | 17.62713938 | 17.93993481 | 19.52713305 | 4.939741755 | 5.876349601 | 4.77826174 |
| ENSMUSG00000063415 | Cyp26b1 | protein_coding | -2.433008093 | 1.275E-70 | 6.906E-69 | 2.807464907 | 2.969748177 | 2.979261223 | 0.521094145 | 0.565196593 | 0.598251027 |
| ENSMUSG00000022206 | Npr3 | protein_coding | -2.44049177 | 3.63E-211 | 7.45E-209 | 12.06362171 | 12.53366837 | 12.5661223 | 2.132737493 | 2.529696369 | 2.468794771 |
| ENSMUSG00000034171 | Faah | protein_coding | -2.505755194 | 3.634E-37 | 1.119E-35 | 3.961526047 | 3.087050308 | 3.220584034 | 0.74047784 | 0.586343567 | 0.564135874 |

**Table S3.** The details for lentivirus information (hanbio, Shanghai, CN)

| | |
|---|---|
| Gene name | Plin1 |
| CDS length | 1554 bp |
| GC content | 50% |
| Species origin | mouse |
| Maker | ZsGreen-PURO |
| Tag | 3 x flag |
| Control virus maker | ZsGreen-PURO |
| Transcript ID | NM 001113471.1 |
| Promoter | CMV |
| Other remakers | The virus carries green fluorescence and piromycin resistance. |

**Gene sequence**

**5’-**ATGTCAATGAACAAGGGCCCAACCCTGCTGGATGGAGACCTCCCTGAGCAGGAGAACGTGCTCCAG
AGAGTTCTGCAGCTGCCTGTGGTGAGCGGGACCTGTGAGTGCTTCCAGAAGACCTACAACAGCACCA
AAGAAGCCCACCCCCTGGTGGCCTCTGTGTGCAATGCCTATGAGAAGGGTGTACAGGGTGCCAGCAAC
CTGGCTGCCTGGAGCATGGAGCCGGTGGTCCGTCGGCTGTCCACCCAGTTCACAGCTGCCAATGAGTT
GGCCTGCAGAGGCCTGGACCACCTGGAGGAAAAGATCCCGGCTCTTCAATACCCTCCAGAAAAGATC
GCCTCTGAACTGAAGGGCACCATCTCTACCCGCCTTCGAAGCGCCAGGAACAGCATCAGTGTGCCCAT
TGCAAGCACCTCTGACAAGGTTCTGGGGGCCACTCTGGCCGGCTGCGAGCTTGCCTTGGGGATGGCCA
AAGAGACAGCAGAATATGCCGCCAACACCCGGGTTGGCCGACTGGCCTCTGGAGGGGCTGATCTGGC
TCTGGGAAGCATCGAGAAGGTGGTAGAGTTCCTCCTGCCACCAGACAAGGAGTCAGCCCCTTCTTCCG
GACGGCAGAGGACCCAGAAGGCTCCCAAGGCCAAACCAAGCCTTGTGAGGAGGGTCAGCACCCTGG
CCAACACTCTTTCTCGACACACCATGCAAACCACAGCATGGGCCCTGAAGCAGGGCCACTCTCTGGCC
ATGTGGATCCCGGGTGTGGCACCCCTGAGCAGCCTGGCCCAGTGGGGCGCATCGGCAGCCATGCAGGT
GGTGTCCCGGCGGCAGAGTGAGGTGCGGGTGCCCTGGCTGCACAACCTGGCAGCCTCTCAGGATGAG
AGCCATGACGACCAGACAGACACAGAGGGAGAGGAGACAGACGACGAGGAGGAGGAAGAAGAGTC

CGAGGCTGAGGAGAACGTGCTCAGAGAGGTTACAGCCCTGCCCAACCCGAGAGGCCTCCTGGGTGGT
GTGGTACACACCGTGCAGAACACTCTCCGGAACACCATCTCCGCAGTGACCTGGGCACCTGCGGCTGT
GCTGGGCACGGTGGGAAGGATCCTGCACCTCACACCAGCCCAGGCTGTCTCCTCTACCAAAGGGAGG
GCCATGTCCCTATCCGATGCCCTGAAGGGTGTTACGGATAACGTGGTAGACACTGTGGTACACTATGTG
CCGCTTCCCAGGCTGTCCCTGATGGAGCCCGAGAGCGAATTCCGAGACATCGATAACCCTTCAGCAGA
GGCGGAGCGCAAAGGGTCCGGGGCGCGGCCCGCCAGCCCGGAGTCCACCCCGCGCCCGGGCCAGCC
CCGCGGCAGCTTGCGCAGCGTGCGGGGTCTCAGCGCGCCCTCCTGCCCCGGCCTGGACGACAAAACC
GAGGCGTCAGCGCGTCCCGGCTTCCTGGCTATGCCCAGAGAGAAGCCTGCGCGCAGAGTCAGCGACA
GCTTCTTCCGGCCCAGCGTCATGGAGCCCATCCTGGGCCGCGCGCAGTACAGCCAGCTGCGCAAGAAG
AGCTGA-**3’**

### 3. Original images of western blotting

**Figure2C** Original WB images in this study including repeats

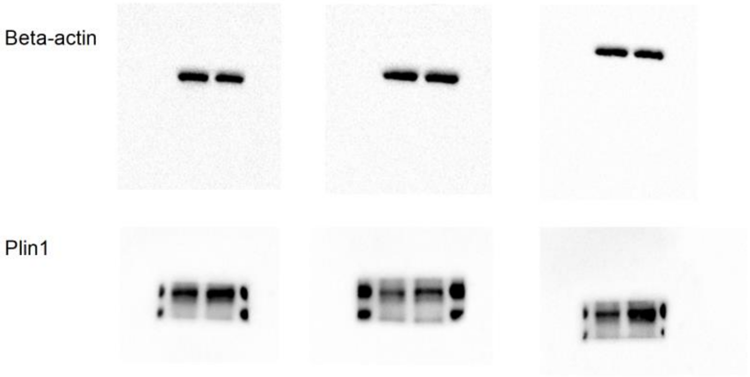


**Figure2N** Original WB images in this study including repeats

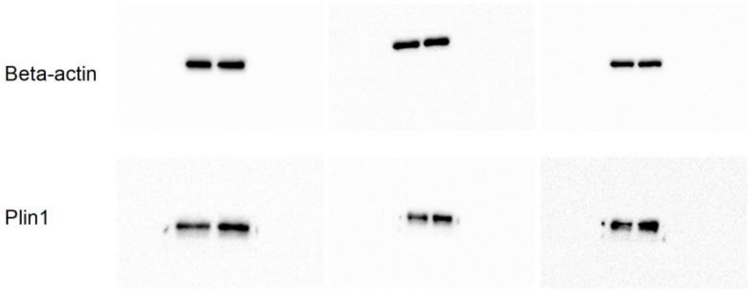


**Figure4A** Original WB images in this study including repeats

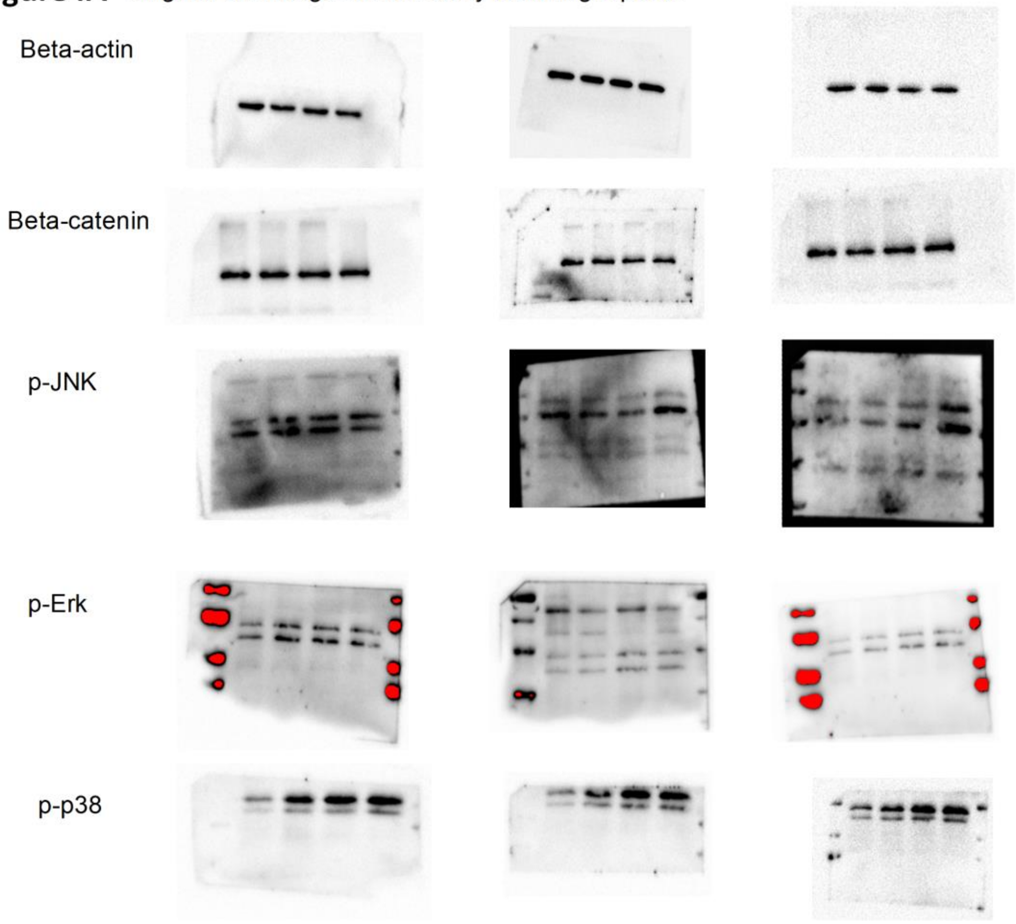

**Figure5A** Original WB images in this study including repeats

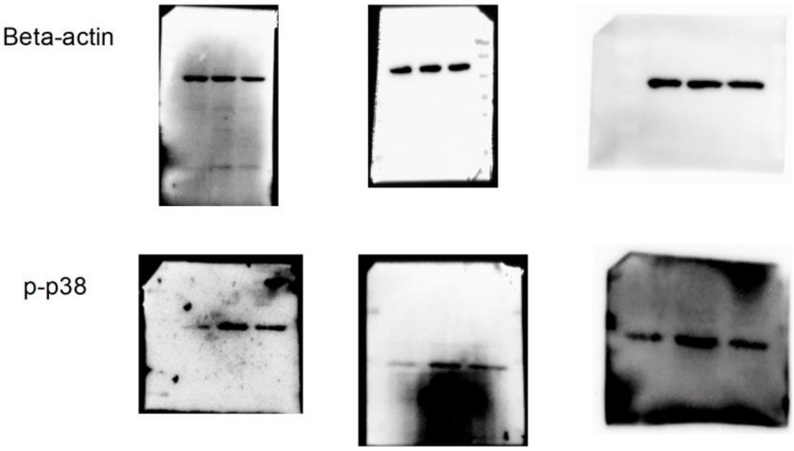


**Figure6A** Original WB images in this study including repeats

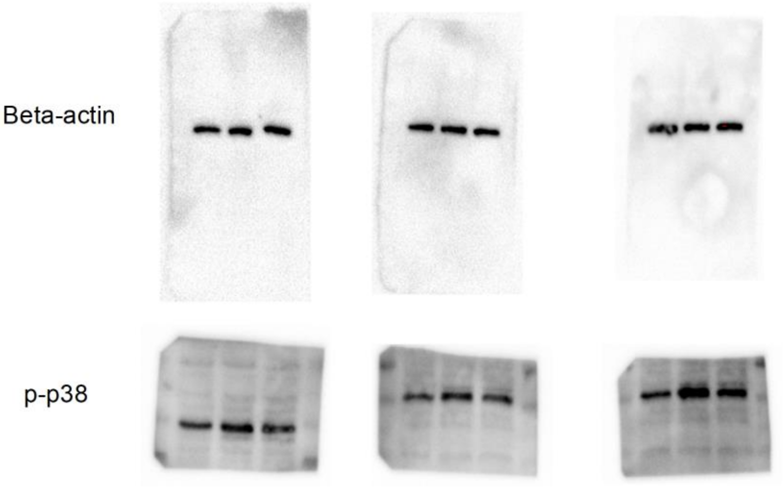
